# Automated muscle path calibration with gradient-specified optimization based on moment arm

**DOI:** 10.1101/2024.05.09.593463

**Authors:** Ziyu Chen, Tingli Hu, Sami Haddadin, David W. Franklin

## Abstract

**Objective:** Muscle path modeling is more than just routing a cable that visually represents the muscle, but rather it defines how moment arms vary with different joint configurations. The muscle moment arm is the factor that translates muscle force into joint moment, and this property has an impact on the accuracy of musculoskeletal simulations. However, it is not easy to calibrate muscle paths based on a desired moment arm, because each path is configured by various parameters while the relations between moment arm and both the parameters and joint configuration are complicated.

**Methods:** We tackle this challenge in the simple fashion of optimization, but with an emphasis on the gradient; when specified in its analytical form, optimization speed and accuracy are improved.

**Results:** We explain in detail how to differentiate the enormous cost function and how our optimization is configured, then we demonstrate the performance of this method by fast and accurate replication of muscle paths from a state-of-the-art shoulder–arm model.

**Conclusion and Significance:** As long as the muscle is represented as a cable wrapping around obstacles, our method overcomes difficulties in path calibration, both for developing generic models and for customizing subject-specific models.

## 1. Introduction

IN musculoskeletal modeling, the geometry of a muscle is often represented by a series of straight and curved cables, namely the muscle path. It is defined by the locations of the origin, via, and insertion points as well as the size(s), location(s), and orientation(s) of the obstacle(s) around which the cables wrap [1]–[4]. Although such a path is a geometrical simplification of the muscle, which alters properties related to the 3-D architecture, it can still have similar muscle length–joint angle and moment arm–joint angle relations to its anatomical reference, provided the path is well configured [5]–[8]. These two relations are crucial to simulation accuracy since muscle length is a major variable in the contraction dynamics [9]–[13] and moment arm directly determines the kinetic capacity of a muscle [14]–[16].

The calibration of a muscle path is not only a fundamental step for the development of a generic musculoskeletal model, but also necessary in subject-specific applications [17]–[21]. Scaling of skeletal geometry, for example, is a common practice of model individualization, where the sizes and shapes of the subject’s segments may be replicated in the model based on anthropometric measurements or inverse kinematics [2], [22]. However, such a scaled model is not yet subject-specific, because the individual characteristics of muscle length and moment arm are only reflected in musculoskeletal geometry, which is different from skeletal geometry [2], [12]. In fact, the scaling of skeletal geometry might distort the biomechanical characteristics which were calibrated to be generically correct. For example, depending on how the path of the triceps surae is defined, enlarging the tibia might result in the Achilles tendon moment arm being increased, unchanged, or even decreased. Thus, for a subject-specific model, a rework of path calibration is necessary after scaling to reflect the individual characteristics of muscle length and moment arm, or at least assure that they remain similar to those in the generic model. Muscle path calibration is performed based on experimental data such as geometric coordinates from medical images [17]–[19], [23] or moment arm measurements [6], [24], and this process can be laborious. To begin with, there is a large number of path-related parameters to be tuned. Each muscle path is defined by at least six parameters, including the two 3-D coordinates of the origin and insertion points. In addition, there will be three extra parameters for each via point and many more when obstacles are included to recreate the anatomical feature of muscle geometry [5], [18]. For example, in the prevalent open-source modeling platform OpenSim [22], cylinders are often used as obstacles to prevent the muscles from penetrating into the joints [7], [24], [25], where the cylinder size, orientation, and location would respectively need one, two, and three parameters to define. In general, monoarticular muscles require between six to 12 parameters, whereas complicated multiarticular muscles may require up to 30 parameters [18].

The size of parameters is not necessarily a big challenge when one knows which to tune. However, if an objective function is too complex, the sensitivity to each parameter is often unclear [21]. Hence, manual tuning tends to be puzzling: e.g., enlarging the obstacle or shifting it away from the center of rotation might increase the corresponding moment arm, but the effect may differ when the joint configuration changes. Even with optimization, the tuning process is not labor-free, if weighting factors are involved and need to be tested repeatedly due to the absence of any knowledge regarding the sensitivity. The second challenge in muscle path calibration lies in the high dimensional relation between moment arm and joint configuration. Each muscle, even monoarticular ones, can actuate multiple degrees of freedom (DoF), and each DoF corresponds to one moment arm [26]–[28]. For instance, the gastrocnemius is a biarticular muscle crossing the knee and ankle, which can be considered to contain at least three DoFs (ankle plantar-/dorsiflexion, ankle eversion/inversion, knee extension/flexion). Therefore, besides the well-studied Achilles tendon moment arm in the sagittal plane [29]–[31], it has an extra ankle moment arm [32], [33] and a moment arm about the knee [26], [34]. With all DoFs considered, path calibration involves matching multiple moment arm–joint angle relations altogether, and it is common to run into trouble where the calibration of one moment arm can distort others that are previously calibrated.

There is more to this challenge than only the high dimensional relation, because each moment arm is also affected by all actuating DoFs of the muscle [32], [35], [36]. Suppose the knee moves while the ankle is immobilized, in theory the Achilles tendon ankle moment arm might still change despite no ankle motion. With this, the moment arm–joint angle relation is no longer depictable by a curve or a surface, but rather requires a hypersurface to demonstrate. For example, the soleus has two moment arms around the ankle, and trying to calibrate them is similar to matching two pairs of surfaces (imagine two heatmaps), which is difficult but viable. However, for the gastrocnemius, each of its three moment arms is dependent on three DoFs, which means that depicting the relation of each moment arm with the DoFs is similar to plotting a volumetric heatmap in 3-D space. In this case, the task of calibration becomes matching the color for three pairs of 3-D heatmaps. While difficult to imagine, manual calibration is theoretically still possible if they are somehow matched slice by slice, where each slice is a 2-D heatmap. But with one more moment arm, e.g. for the rectus femoris or many of the shoulder muscles, the number of dimensions to be matched is beyond three, forcing manual tuning to be simplified and compromising the overall model accuracy.

In light of these challenges, it takes extensive effort to develop musculoskeletal models to simulate many different motions or to individualize them for each subject. This difficulty hinders the advancement and application of musculoskeletal modeling and simulation. Therefore, here we develop a gradient-based method for automated muscle path calibration. Our goal is to tune path-related parameters so that the moment arm–joint angle relation of a model matches with the target specification. An optimization framework is established for a classic muscle path wrapping method [1] and the gradient for the cost function is derived in its analytical form to increase optimization speed and accuracy. The concept employed in the gradient derivation is universal and may be applied to the model calibration of many other complex systems.

## II. Methods

### A. Optimization Method

The process of parameter tuning for a muscle path is formulated as a least-squares problem with the cost function

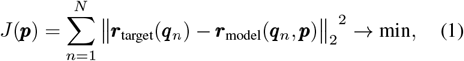

where ***p*** denotes muscle path parameters, and ***r***(***q***) is the moment arm at joint configuration ***q***. Note that ***r*** and ***q*** are vectors with the dimension of DoF number, whereas *N* is the number of joint configurations. The target value and the model output are indicated by the subscripts, but to make the notations concise, the model output will also be abbreviated as ***r***(***q, p***), ***r***(•, ***p***), or ***r***.

In this study, we limited the obstacle type to cylinder to simplify the discussion, but the same principle applies to other types such as sphere and sphere-capped cylinder described in [1]. Currently, our optimization method requires the composition of each muscle path to be decided in advance, which is based on three fundamental segment types: straight (no obstacle), single-cylinder, and double-cylinder. A path can be constructed as an individual segment or a combination of multiple different segments, e.g.,

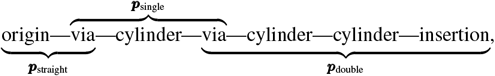

and the composition of ***p*** is correspondingly based on three fundamental forms:

- ***p***_straight_: (^j^***u***_P_, ^j^***u***_S_)
- ***p***_single_: (^j^***u***_P_, ^j^***u***_S_, *R*, ^j^***u***_C_, *α, β*)
- 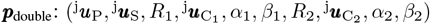

where ^j^***u***_P_, ^j^***u***_S_ ∈ ℝ^3^ are the coordinates of the anchor points of a path segment expressed in their reference frames, written as column vectors; e.g., ^j^***u***_P_ = [^j^*x*_P_ ^j^*y*_P_ ^j^*z*_P_]^T^. The radius of a cylinder is denoted by *R* ∈ ℝ^+^ and the center by ^j^***u***_C_ ∈ ℝ^3^, whereas the orientation is defined by two Euler angles *α, β* ∈ ℝ. In the following discussion, the start and end anchor points of a path segment are denoted as P and S respectively, and when wrapped by the cylinder (whose center is denoted as C), the two wrapping points are denoted as Q and T (Fig. 1). Also, the frame in which coordinates are expressed are indicated by the superscript: with w for the world frame, c for the cylinder frame, and j for one of the joint frames; see Appendix A for coordinate transformation.

**Fig. 1.**
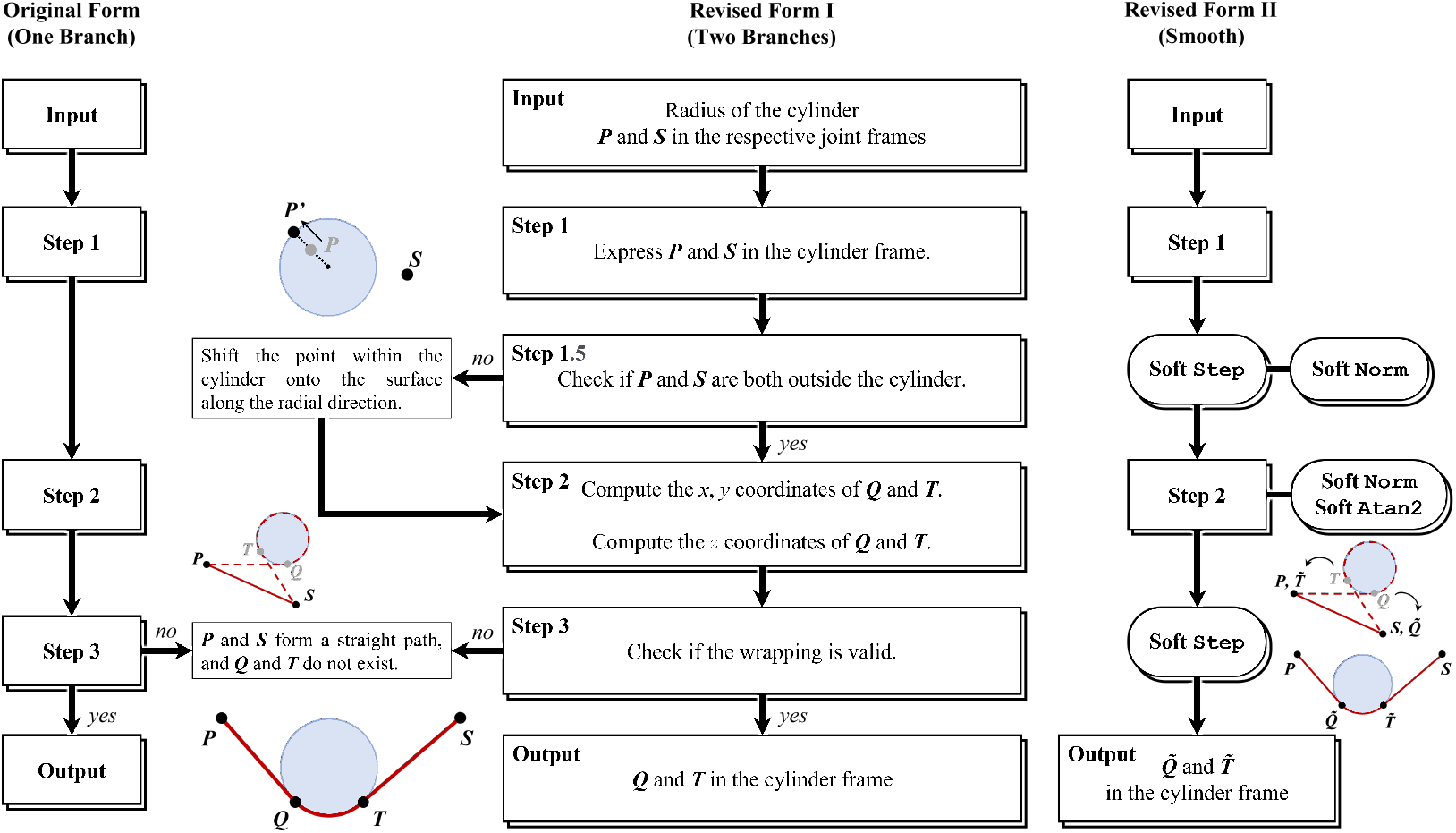
Workflows of the original and revised obstacle-set methods. Left: the original form with one conditional statement. Middle: the revised form with an extra conditional statement for continuity. Right: the smooth form revised for optimization. The naming and order of the anchor and wrapping points in the single-cylinder case are indicated by the illustration in the bottom.

A muscle path is configured by ***p*** using the obstacle-set method [1], which computes the potential wrapping points on the obstacle(s) by finding the minimum-distance path between the two anchor points in each segment. To reduce computational load, we first took a geometric approach to compute ***r***, which is based on the principle of virtual work and is algorithmically efficient by computing the velocity terms using the Kane’s method [37]–[39]. Then, since the typical algorithms solving (1) (e.g., Levenberg-Marquardt or trustregion) require the gradient *∂J/∂****p***, and by the chain rule

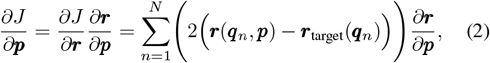

we need to specify *∂****r****/∂****q*** in its analytical form.

Notice for a path composed of *I* segments, its length (*l*) is the sum of the lengths of all segments, and the same applies to its moment arm (*∂l/∂****q***). This means (2) can be decomposed as individual computations for each path segment with

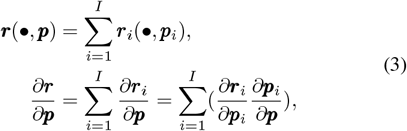

where the *i*-th segment is configured by ***p***_*i*_, either in the form of ***p***_straight_, ***p***_single_, or ***p***_double_. Therefore, essentially our goal is to specify *∂****r***_*i*_*/∂****p***_*i*_, and knowing that the case of any multisegment path can be broken down to an individual segment, our following discussion of moment arm focuses on the path segment and we drop the subscript *i* for convenience.

Direct derivation of *∂****r****/∂****p*** is overwhelming considering its structural complexity. It would require a tremendous amount of effort, and the eventual result would be filled with pages of repeated terms and computationally inefficient. Thus, we circumvented this problem by disassembling it into a composite of multiple gradients whose analytical forms are easy to derive. For instance, moment arm can be computed given the anchor and wrapping points (as in ***r***(***u***)), and by the chain rule

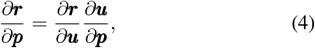

where ***u***(***p***) is obtained with the obstacle-set method. As we will show later, ***r***(***u***) is relatively simple in structure, so *∂****r****/∂****u*** is not difficult to derive, and our remaining goal is to compute *∂u/∂****p***. However, ***u***(***p***) is still complex and contains indifferentiable components, so we began with revising the obstacle-set method into a smooth form, such that *∂u/∂****p*** exists for all ***p***. This is accomplished by replacing the conditional statements in the obstacle-set method as well as other nondifferentiable components with soft functions which yield almost the same outputs but are continuously differentiable (Fig. 1).

### B. Soft Functions

Fig. 1 (middle) shows the detailed workflow of the revised obstacle-set method, containing two conditional statements. The first statement checks if either of the anchor points is inside the cylinder. Note that this was not included in the original method by [1] (Fig. 1, left), and we introduced it to simplify the optimization since it removes complex geometric constraints between the anchor points and the obstacle. The second statement checks if the wrapping angle exceeds 180°, which is considered invalid wrapping. Naturally, conditional statements make ***r***(*•*, ***p***) nondifferentiable in certain domains, and we introduced soft functions to patch them (Fig. 1, right).

Step 1.5 essentially keeps a minimum distance of *R* between the point and the cylindrical axis (e.g., 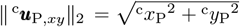 to prevent Step 2 from returning Inf or NaN. In our case, if for example P locates inside the cylinder, it is shifted onto the surface along the radial direction as P^*′*^ (Fig. 1, middle), and the coordinates become

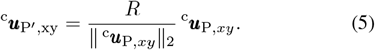

More specifically,

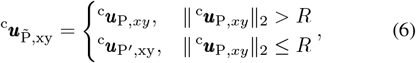

and the coordinates of such transitional point calculated using a soft version of Heaviside step function:

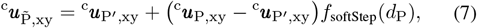

where *d*_P_ = ^c^*x*_P_^2^ + ^c^*y*_P_^2^ *− R*^2^, and

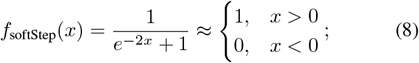

the input can be magnified to steepen the transition around 0. Now, the conditional statement of (6) is smoothed: Premain as P if *d*_P_ *>* 0, otherwise it approaches the shifted point P^*′*^ when *d*_P_ tends to 0 in the negative direction.

With 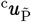 and 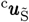, Step 2 finds the appropriate points of tangency Q and T on the cylinder (^c^***u***_Q_ and ^c^***u***_T_; see (S11) and (S13) in Appendix C). Then based on the sign of

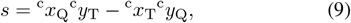

Step 3 returns either the coordinates of the two wrapping points or null if wrapping is invalid (Fig. 1, left). The latter situation can be problematic as it leads to yet another conditional statement in moment arm computation, in which different formulas are used depending on whether the wrapping occurs:

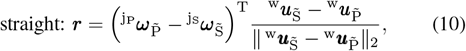

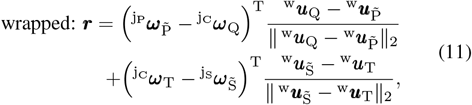

where ***ω*** is obtained using (S2) in Appendix A.

Here, the revision of Step 3 into a smooth form involves a transition from (10) to (11) when *s* changes from positive to negative, and the complication arises with the involvement of joint frames: P, S, and C could be fixed on separate bones; i.e., the implication of j_P_, j_S_, and j_C_ may be different. This makes it hard to cancel out the terms in (11) to transition into (10). To this end, we have Q and T respectively approaching S and P when *s* tends to 0 in the negative direction (Fig. 1, right), and similarly this can be achieved with (8):

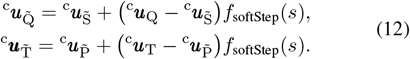

This way, moment arm computation is generalized as

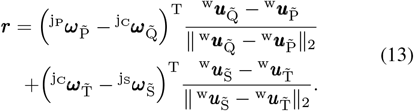

When 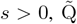 and 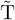 maintain their original coordinates as wrapping points Q and T (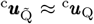and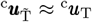), and moment arm is computed with (11). When 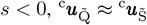 and 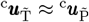, and (13) simplifies into (10). Importantly, this generalization accommodates the wrapping and moment arm computation involving two obstacles.

Furthermore, Step 2 involves the calculation of the 2-norm (also in (5) and (13)) and the 2-argument arctangent (see (S13) in Appendix C). Both are nondifferentiable at the origin, and we replaced them with

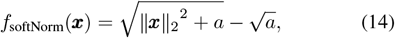

and

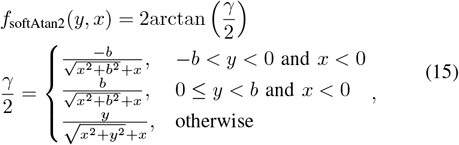

where *a* and *b* should be sufficiently small positive numbers (e.g., `.eps` and 0.001 respectively).

In such a manner, we have modified a branched workflow into a smooth form that is differentiable across the entire domain of its input parameters. This modification is verified using kinematic measurements to ensure that its difference with the non-smooth form is small enough across a large range of motion (see Appendix B).

With (8), the obstacle-set method (i.e., the computation of ***u***(***p***)) is revised into a smooth form, and since (8), (14), and (15) are clearly all differentiable, it is now possible to compute the gradient *∂u/∂****p***. As shown in (4), our target function ***r***(*•*, ***p***) is a composite of ***u***(***p***) and the generalized moment arm computation ***r***(•, ***u***) with (13). However at this stage, *∂****r****/∂****p*** is still difficult to compute due to the knotty structure of *∂u/∂****p***, so we continue down the chain; see Appendix C for the mathematical derivation in detail.

### C. Calibration and Validation

With the gradient *∂****r****/∂****p*** specified, we used the nonlinear least-squares solver (lsqnonlin) in MATLAB to solve (1). The computation of this cost function requires a musculoskeletal model and the input of some kinematic dataset ***q*** for the model output, and the correspondent moment arm data at ***q*** are required as the target value. Ideally, ***q*** should cover joint configurations as diverse as possible for accurate calibration. For the model output, we used a 12-DoF 42-muscle human shoulder–arm model [40] (with muscle path from [18]; see all DoFs and muscle paths in Appendix D), and the kinematic measurements of the shoulder and arm from 10 intransitive daily tasks performed by a single subject in [41]. The kinematics from five tasks (gesturing an OK sign, pumping fists, blocking out light from the face, saluting, and pointing) were input to the reference model to generate artificial calibration data, while ***q*** from another five tasks (gesturing a thumb-down, signaling for hitchhike, greeting, gesturing to stop, and gesturing for silence) were used for validation.

For the target value, we utilized the same model as reference to generate artificial moment arm data; that is, (1) becomes

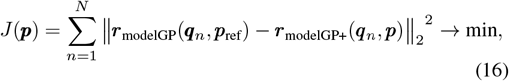

where ***r***_modelGP_ is computed based on the slightly revised obstacle-set method by [1] (Fig. 1, middle) with ***p***_ref_ from [18], while ***r***_modelGP+_ is based on our smoothed form (Fig. 1, right). Thus in a sense, this process is equivalent to replicating the muscle path geometry in the reference model. The input of ***q*** was interpolated from the aforementioned kinematic measurements. This is because the original dataset consists of joint configurations from a massive number of time instants (1000-Hz sampling rate), many configurations are similar and hence redundant as input for producing the target value. For calibration, to avoid an underdetermined system, the number of joint configurations equals to the number of path parameters that each muscle has (Supplementary Table II); e.g., an 18-parameter path will be calibrated based on the kinematics in a total of 18 instants from five movements (*N* as 18 in (16)). For validation, five instants were extracted from each movement, that is 25 joint configurations in total.

With the model output and target value in place, we set out to demonstrate the performance of our method through a comparison test as well as the implementation of muscle path calibration. In the test, the solver was configured to run with and without the gradient specified analytically, and we compared the optimization results for some representative muscle paths. In order to take algorithmic features into account, the comparison between gradients was repeated for two `lsqnonlin` algorithms. In the implementation, the solver was tasked to calibrate the moment arms of all 42 muscle paths, and we manipulated the input using four variants representing progressive levels of application complexity. The input variants were generated based on a 2×2 design: original/modified parameter structure × noise-free/noisy calibration data. For both the test and the implementation, optimization was performed on a 2.9GHz Intel Core i9 with 64 GB RAM and 14 CPU cores using parallel computing (`parfor`): The processor was not overclocked, and the number of parallel workers was set as 14. We structured parallel computing to optimize 14 initial points simultaneously for a single muscle path within each `parfor` loop. Other details are described as follows.

The comparison test aims to evaluate the contribution of gradient specification as well as to determine an appropriate configuration for the implementation. First, we tested the `levenberg-marquardt` algorithm on two single cylinder–based (serratus anterior, superior and middle parts) and two double cylinder–based muscle paths (latissimus dorsi, thoracic and iliac parts), each with 70 sets of initial points. The `levenberg-marquardt` algorithm is relatively simple in that it has only one type of search direction [42], [43], and the optimization process should generally be similar when the analytical and numerical gradients differ only by numerical error. So with this test, we may also verify if our computation of the analytical gradient is correct. The initial points for each path—including locations (mm), radius (mm) and Euler angles (rad)—were randomly generated using `rand` in five ranges (from [−1, 1] to [−10^4^, 10^4^] with an increment of magnitude in between) and optimized without any constraints for our method to be verified across a vast domain. Then, we tested `trust-region-reflective,` which is the default algorithm for `lsqnonlin` and more robust. Optimization was performed with two additional complex paths (extensors carpi radialis brevis and ulnaris), each with 84 sets of initial points. They were also randomly generated but were bounded with slight anatomical constraints— the same initialization that will be explained later for the implementation. For both tests, `FunctionTolerance, OptimalityTolerance,` and `StepTolerance` were configured as 2.5 ×10^*−*16^ with MaxIterations as 10^3^ and `MaxFunctionEvaluations` as 10^5^, and the numerical gradient was computed using MATLAB’s built-in forward-difference method.

Based on the preliminary results, the implementation of calibrating 42 muscle paths was configured as follows:

1. `trust-region-reflective` (gradient-specified);
2. 10^*−*4^ for `StepTolerance,` and 2.5 *×* 10^*−*16^ for `FunctionTolerance OptimalityTolerance;`
3. 10^2^ for `MaxIterations` and `Inf for MaxFunctionEvaluations,` and the initial points were generated with the subsequent considerations.

As previously mentioned, each muscle path requires predetermination of the number of via points and cylinders for the structure of ***p***. Some muscle paths are modeled with via points and more than two obstacles (Supplementary Table II), and their structures of ***p*** will be a combination of the three typical forms. For example, the extensor carpi radialis longus is modeled in the form of

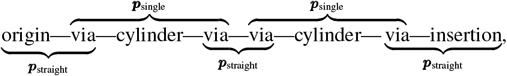

which contains a total of 30 path parameters. To test the robustness of our method against parameter structure, we considered two scenarios. First, we set the structure of path parameters to the same as in the reference model. This is mostly the case of subject-specific modeling, where the parameter structure is known in the generic model and calibration is performed to adjust parameter values. In the second scenario, all muscle paths were configured with either one or two cylinders and without any via points, resulting in a different parameter structure from their reference counterpart (Supplementary Table II) and approximately 10% fewer parameters in total (***p*** and ***p***_ref_ in (16) may differ in dimension). In other words, with at most two obstacles to wrap around, this path needs to imitate the geometry of the original path, potentially configured by multiple points and obstacles. This is similar to the case of generic modeling, where often one obstacle per joint is placed regardless of how complex the muscle geometry might be.

Apart from the predetermination of parameter structure, We bounded the joint-frame coordinates of the origin and insertion points (not all anchor points) within a uniformly distributed range of ±30 mm from the respective original values in the reference model—in both initialization and optimization. This is not a prerequisite, but since the origin and insertion points attach to anatomical landmarks with accurate measurements, we included this constraint to speed up the calibration. Also, when generating the initial points, the joint-frame coordinates of cylinder center(s) were randomly initialized around the sections of the line between the initial origin and insertion points. This also accelerates the process, since if a cylinder is not wrapped at the beginning of the optimization, *∂****r****/∂* ^j^***u***_C_ remains a null matrix, and ^j^***u***_C_ might be left untuned till the end. For efficiency, we ensured that the cylinders are not too departed from the initial muscle paths.

To reduce the risk of local minimum, the optimization was globally iterated with multiple randomly generated initial points. For each path, the number of initial points is 14 times the number of parameters (Supplementary Table II); e.g., a 12-parameter path will be calibrated with at most 168 sets of different initial values. Additionally, to realize a trade-off between speed and accuracy, we programmed the calibration to first iterate over 42 initial points (i.e., three `parfor` loops), and if the cost (16) per moment arm per joint configuration does not reach a sufficiently small level *k* (e.g., 0.01), another 42 will be iterated. The rest of initial points will only be iterated when the normalized cost fails to reach 100*k* after the first 84 global iterations, and the calibration stops whenever 100*k* is reached or when all initial points run out. If none of the global iterations reaches the desired threshold, 14 sets of initial points that led to the lowest costs but had the solver exit due to the iteration limit will be further optimized with `MaxIterations` as 10^3^.

Optimization performance is also influenced by the calibration data, thus to examine the robustness of our method against error, the calibration was also performed with noisy data. For this, the aforementioned artificial moment arm data were added to with a composite of relative error (uniformly distributed within ±20% of the reference value) and absolute error (uniformly distributed within ±2 mm). This leads to a total of four conditions for the implementation (original/modified parameter structure × noise-free/noisy calibration data), and each was simulated five times with different sets of initial points to evaluate the performance in terms of calibration speed and validation accuracy. Note that the five sets of initial points were kept the same for both noise-free and noisy conditions, and stopping criteria remained unchanged for all conditions; otherwise, the difference in results might also be attributed to initialization and configuration.

The calibration results are validated by mean absolute error

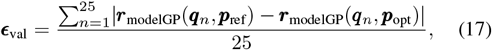

where ***p***_opt_ is the solution of (16). Here, notice the optimized parameters are input into the non-smooth model ***r***_modelGP_ despite obtainment from the smooth form, making the evaluation standard more strict.

## III. Results

We selected a few representative muscle paths to test the optimization with two classic `lsqnonlin` algorithms as well as to compare the performance with and without gradient specification. Using the `levenberg-marquardt` algorithm, the optimization processes based on the analytical and numerical gradients are generally the same, with costs descending in identical fashions for most of the initial points (Fig. 2 and Supplementary Figs. 3–5). Whereas with `trust-region-reflective,` the cost descents much faster with gradient-specified optimization (Fig. 3 and Supplementary Figs. 6–10): On average, by the 100th or even the 10th iteration, the cost minimized based on the analytical gradient is already lower than the final cost achieved by the numerical gradient.

**Fig. 2.**
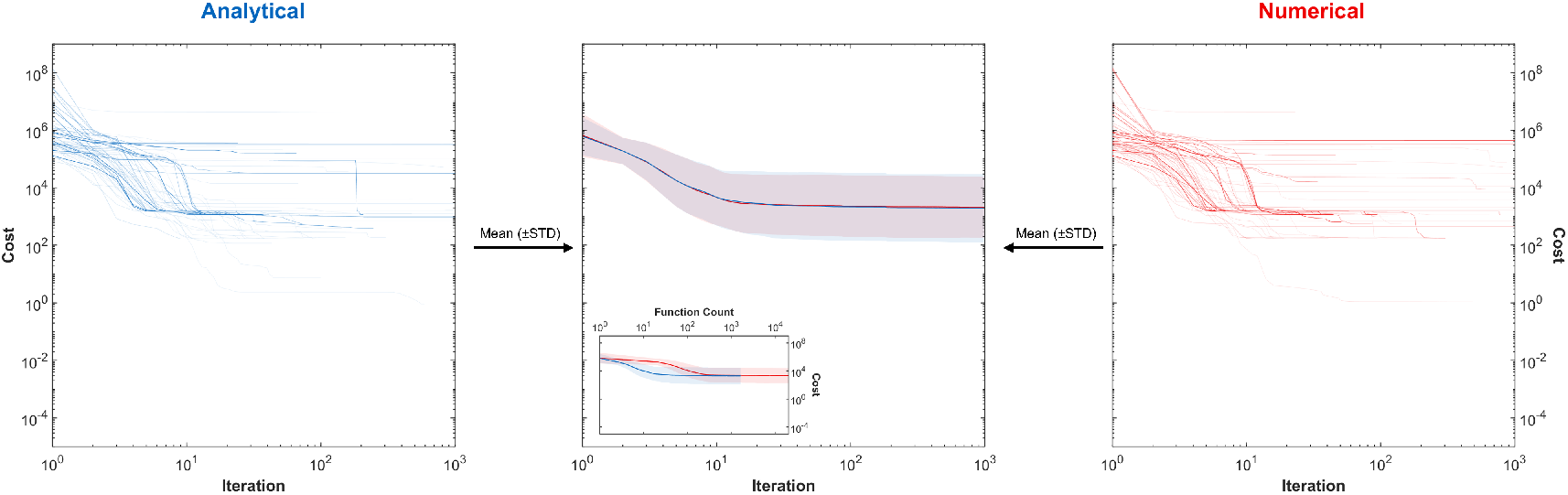
Cost–iteration curves for the optimization using the `levenberg-marquardt` algorithm (latissimus dorsi, iliac part). The results of each initial point from the analytical (blue) and numerical (red) gradients are plotted respectively on the left and right. The mean and STD of the costs are plotted in the middle, with function count–dependent curves in the corner.

**Fig. 3.**
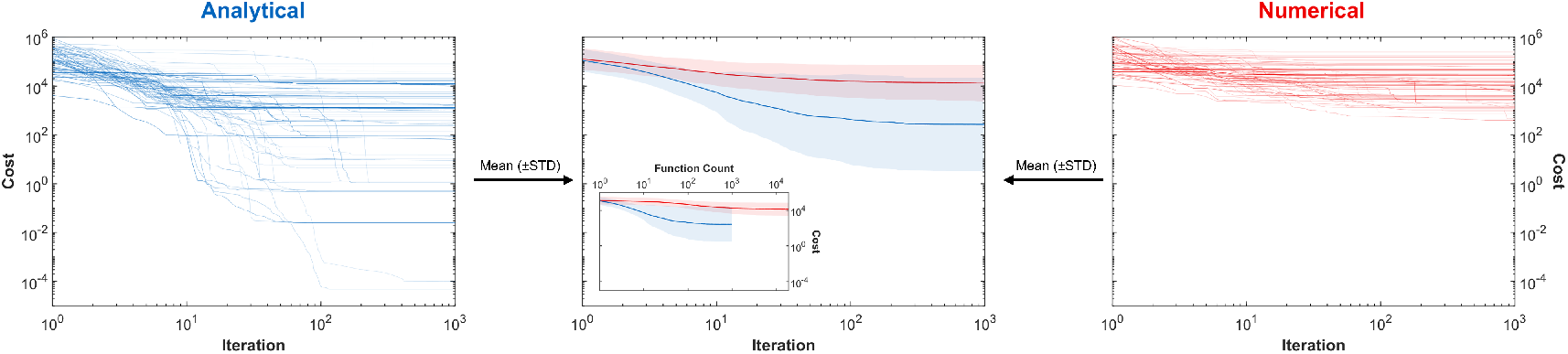
Cost–iteration curves for the optimization using the `trust-region-reflective` algorithm (latissimus dorsi, iliac part). The results of each initial point from the analytical (blue) and numerical (red) gradients are plotted respectively on the left and right. The mean and STD of the costs are plotted in the middle, with function count–dependent curves in the corner.

We also recorded the computation time and the number of function counts (how many times the cost function is evaluated) for each iteration. Fig. 4 shows the ratio of computation time per iteration between the gradient-specified and unspecified optimizations, and as can be expected based on the computational principle of the numerical gradient, this ratio is almost linear to the number of parameters in the cost function. With this ratio, we scaled the cost descent to the number of function counts in Figs. 2 and 3, and Supplementary Figs. 3–10 to demonstrate optimization performance with respect to computation load. From this perspective, with `levenberg-marquardt,` even if the cost reduction per iteration is almost the same between the gradient-specified and unspecified optimizations, the former requires fewer function evaluations and is hence more efficient. As for the `trust-region-reflective` algorithm, the advantage achieved by the analytical gradient is further magnified by this computational efficiency.

**Fig. 4.**
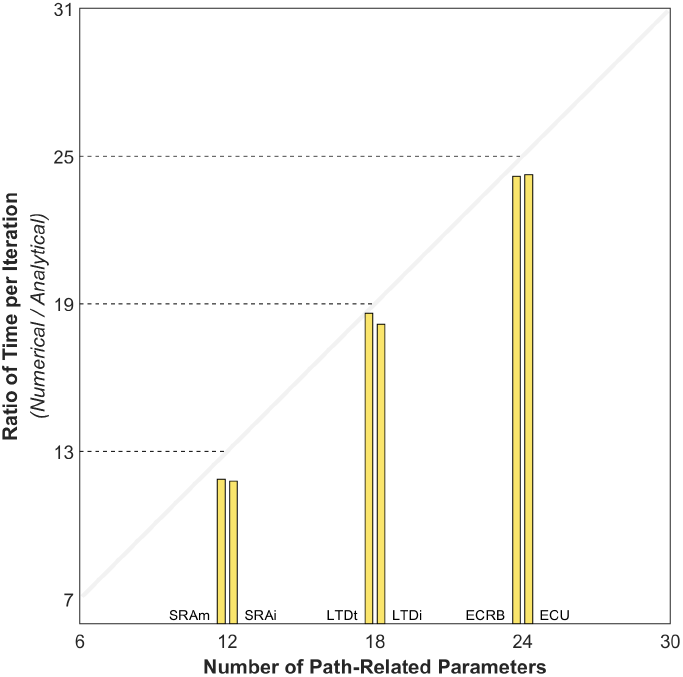
Ratio of computation time per iteration (numerical vs. analytical) of six representative muscle paths. See Supplementary Table II for abbreviations.

To demonstrate the performance of our method in the implementation, we devised four conditions in which 42 muscle paths are calibrated of their multi-dimensional moment arms, and the results are summarized in Table I and visualized in Fig 5 and Supplementary Figs. 11–13. In an ideal condition where the parameters-to-optimize share the same structure with their reference and the calibration data contain no noise, our method completed the calibration task in 7.7 min with parallel computing. Across all 182 effective moment arms, the median validation error is 5 × 10^*−*5^ mm and the max is 3.37 mm for the palmarflexion/dorsiflexion moment arm of the flexor carpi radialis.

**TABLE 1.**
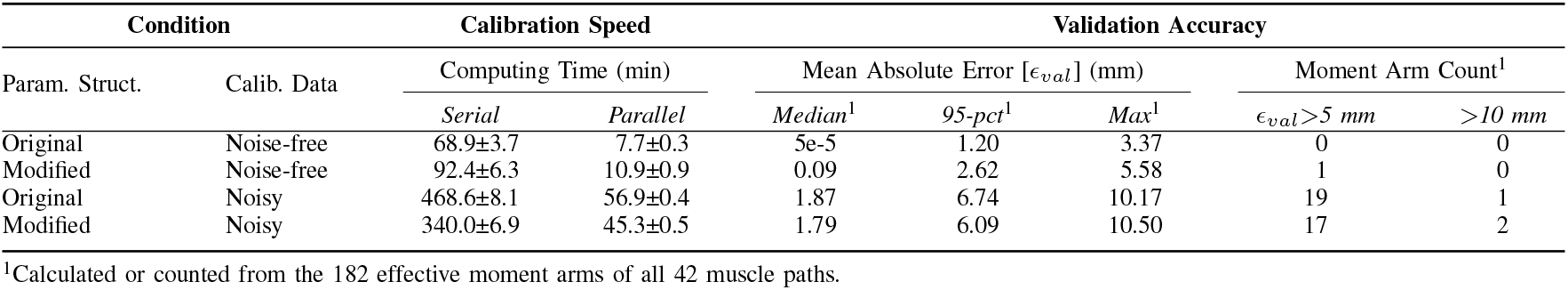
Method performance.

| Param. Struct. | Condition | Calibration Speed |  | Validation Accuracy |  |  |  |  |
| --- | --- | --- | --- | --- | --- | --- | --- | --- |
| | | Computing Time (min) | | Mean Absolute Error [$\epsilon_{val}$] (mm) | | | Moment Arm Count<sup>1</sup> | |
| | | Serial | Parallel | Median<sup>1</sup> | 95-pct<sup>1</sup> | Max<sup>1</sup> | $\epsilon_{val} > 5$ mm | $> 10$ mm |
| Original | Noise-free | 68.9±3.7 | 7.7±0.3 | 5e-5 | 1.20 | 3.37 | 0 | 0 |
| Modified | Noise-free | 92.4±6.3 | 10.9±0.9 | 0.09 | 2.62 | 5.58 | 1 | 0 |
| Original | Noisy | 468.6±8.1 | 56.9±0.4 | 1.87 | 6.74 | 10.17 | 19 | 1 |
| Modified | Noisy | 340.0±6.9 | 45.3±0.5 | 1.79 | 6.09 | 10.50 | 17 | 2 |

<sup>1</sup>Calculated or counted from the 182 effective moment arms of all 42 muscle paths.

**Fig. 5.**
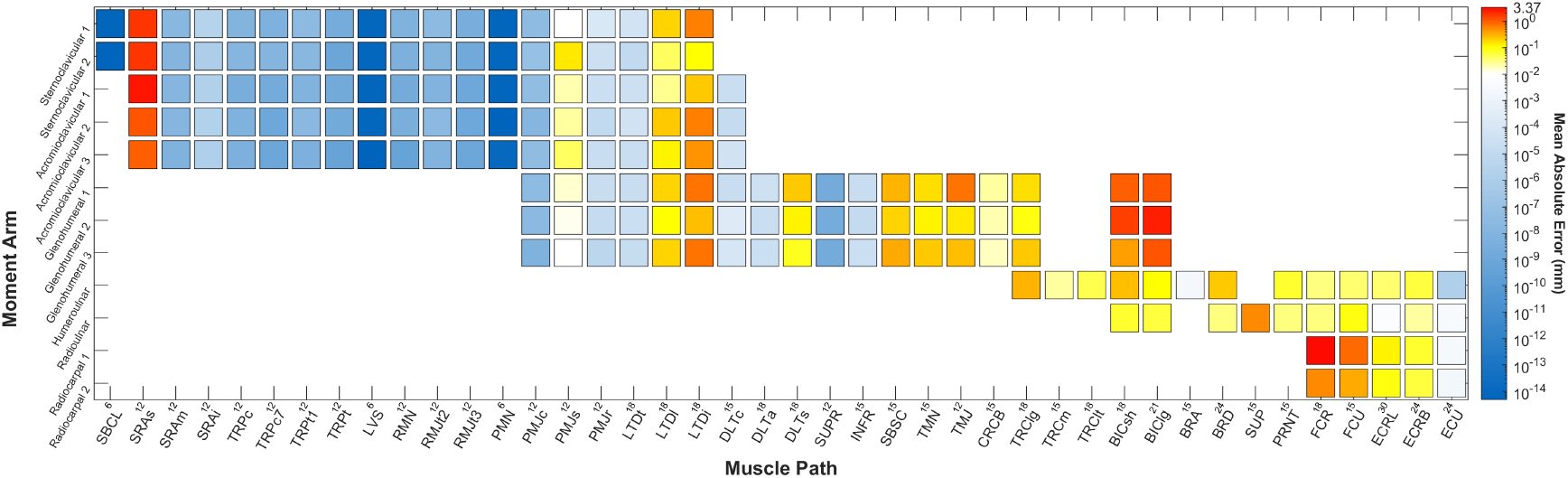
Absolute errors in 182 effective moment arms of 42 muscle paths in validation (original parameter structure and noise-free calibration data). The magnitude of the error is indicated by the intensity of the color. The number in the horizontal axis corresponds to the parameter number for each muscle path. See Appendix D for nomenclatures in the axes.

When the input parameters are modified of their structures (e.g., a path defined by two cylinders is now calibrated with only one cylinder, or a path with multiple via points is now calibrated without them), the calibration time is 10.9 min, and only one moment arm contains error more than 5 mm in validation. When the calibration data are noisy, the performance of our method is reduced, which is not surprising since the introduced noise is quite large (a composite of ±20% relative error and ±2 mm absolute error). Nevertheless, its speed and accuracy are still satisfactory. For example, in the case of the original parameter structure, the calibration task was completed in 56.9 min with only one moment arm containing validation error over 10 mm.

For a more intuitive understanding of our method’s working process, we demonstrate in Figs. 6 and 7 two cases of muscle path calibration, each with a featured complication. In Fig. 6, the initial anchor points are both within the cylinder. This is a situation that the original obstacle-set method cannot handle, and we patched the issue by shifting such a point onto to the cylinder surface to prevent a breakdown in wrapping point computation. As soon as the optimization starts, they are gradually expelled out of the cylinder to reduce the cost, achieving a smooth transition from one branch to another within a conditional statement. Fig. 7 shows a typical path configured by two cylinders, and the misplacement of either cylinder will induce error in moment arm computation. Driven by the gradient, first the orientations of both cylinders are rotated to the correct directions, and then their radii are adjusted to the right sizes—all in tens of iterations.

**Fig. 6.**
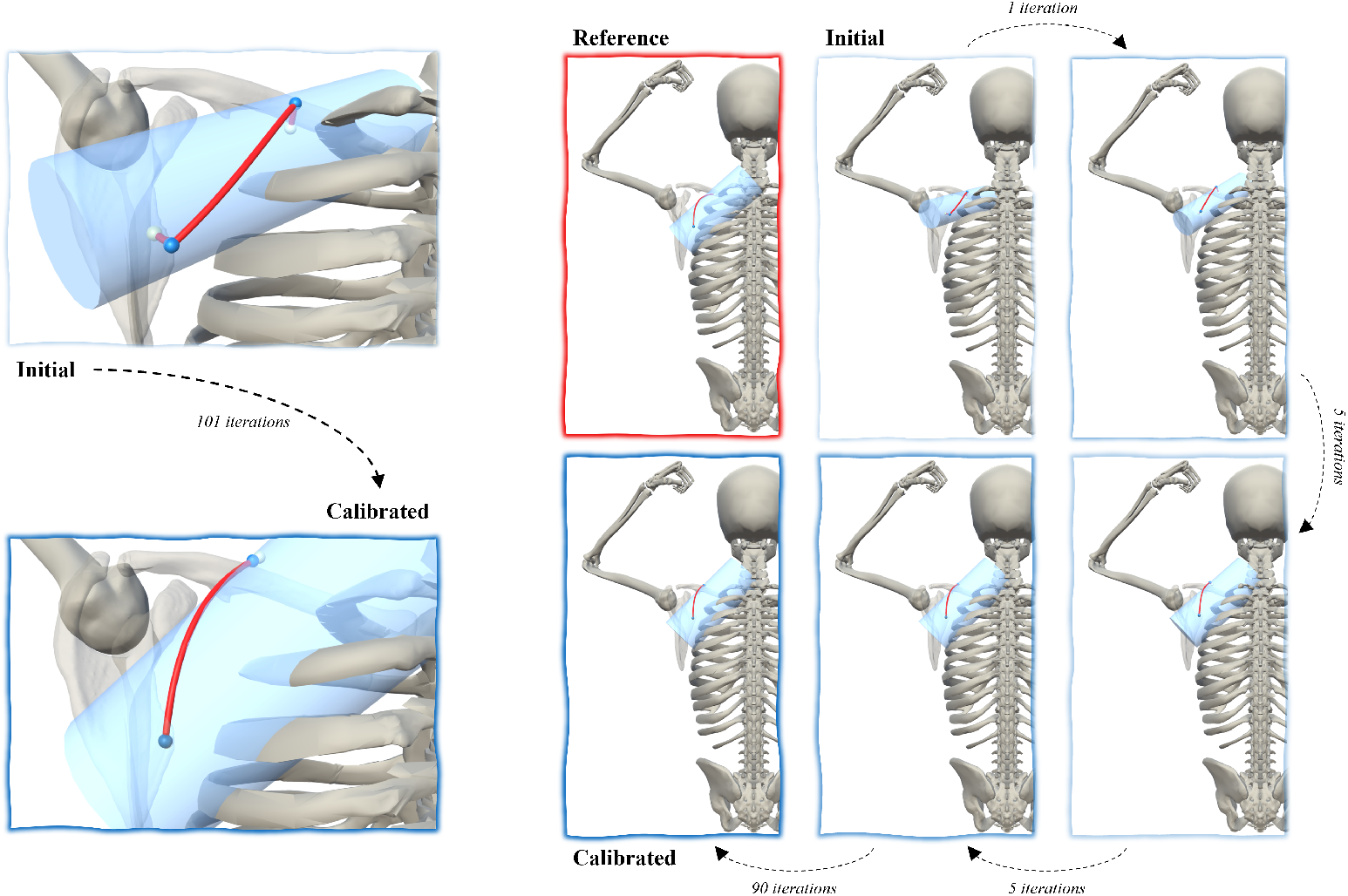
A featured case of path calibration for the serratus anterior (superior part). Left: the anchor points and the obstacle defined by pre- and post-optimized parameters and the resultant wrapping points. Right: the reference path configuration and the optimization process from the initial point to the solution. The yellow points on the red muscle path are the anchor points, while the wrapping points are in blue. The initial anchor points are inside the cylinder, which would induce a computation breakdown in the original obstacle-set algorithm but is patched by our revision. As the optimization progresses, the anchor points are gradually expelled from the cylinder, leading to a path configuration valid for the original algorithm.

**Fig. 7.**
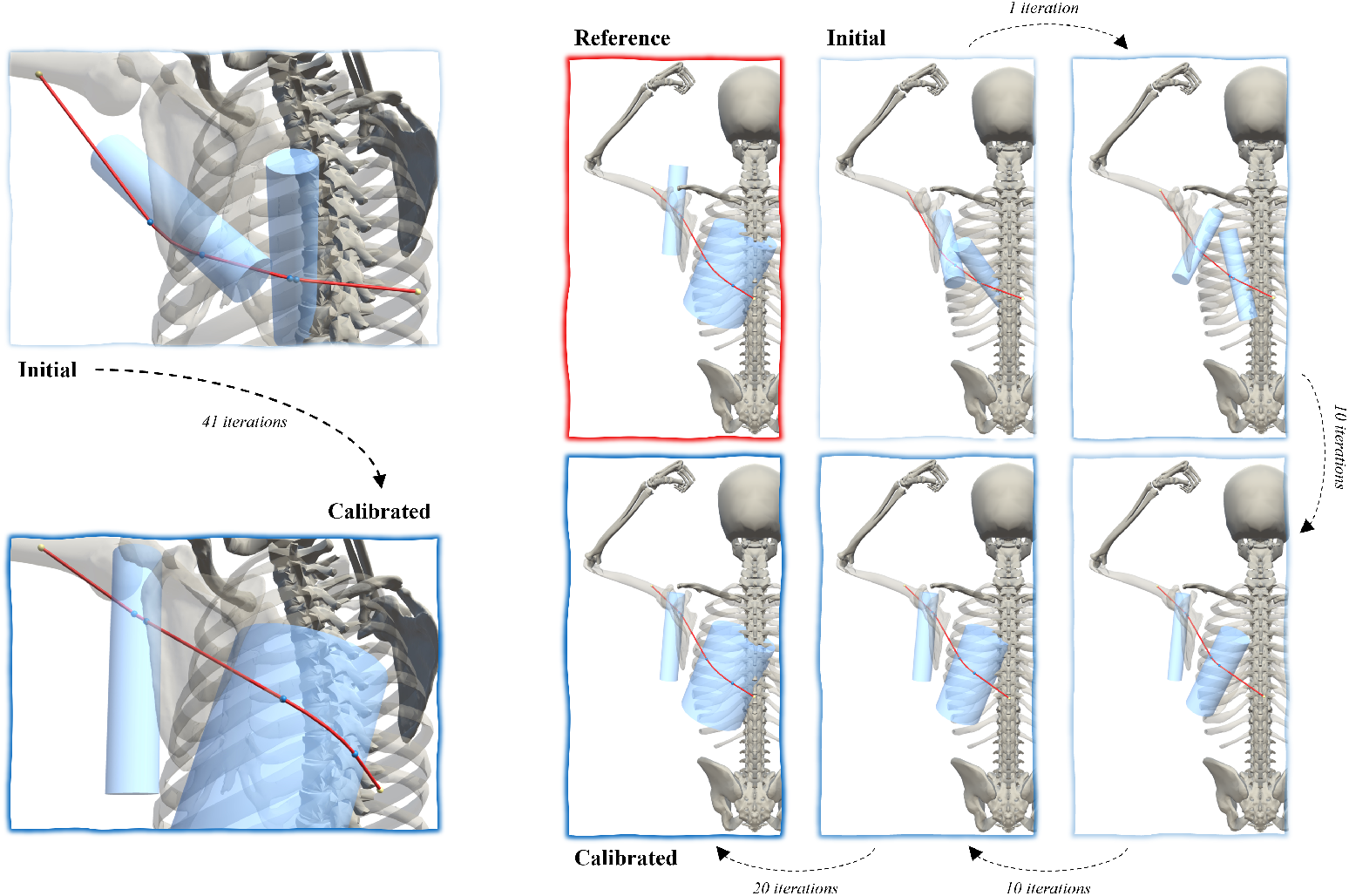
A featured case of path calibration for the latissimus dorsi (thoracic part). Left: the anchor points and the obstacles defined by pre- and post-optimized parameters and the resultant wrapping points. Right: the reference path configuration and the optimization process from the initial point to the solution. The yellow points on the red muscle path are the anchor points, while the wrapping points are in blue. The path is complicated in that it wraps around two obstacles, both of which need to be correctly placed for moment arm calculation to be accurate. By the 11th iteration, the cylinders are oriented to the correct directions, and their radii are adjusted correctly in another 30 iterations. The parameters related to cylinder placement are optimized without constraints.

## IV. Discussion

A key to optimization speed and accuracy is the gradient, which in our case is the gradient of the moment arm function with path-related parameters as variables. To derive it analytically, the cost function must be differentiable in the first place. For this, we revised the obstacle-set method for muscle path wrapping by substituting its conditional statements and nondifferentiable components with soft functions. Next, we disassembled the revised function into smaller modules that are easier to derive separately, and then assembled the gradients back into the desired gradient composite. The result is simplistic in form and thus easy to compute, which further increases optimization speed. Importantly, the concept of the chain rule is universal and may be useful for deriving the gradient of other complex functions as well.

Specifying the gradient in its analytical form is more than a formal alternative to gradient approximation using the typical finite difference method. As we demonstrate in our preliminary tests, the performance of the gradient-specified optimization is distinct from that of the gradient-unspecified. This arises from how the analytical and numerical gradients are computed as well as how each optimization algorithm proceeds with different gradients. For example, to estimate the gradient numerically with the forward-difference method, consider

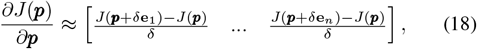

where **e**_*i*_ is the standard basis vector with the *i*-th element as 1 and 0s elsewhere, and *δ* is a sufficiently small positive value. To approximate the gradient, the additional number of function evaluations equals to the dimension of input parameters; this number will double if a central-difference method is used. In other words, the time complexity of (18) is *O* (dim(***p***)) [44]–[46]. However, if the analytical form is explicit, the cost function only needs to be evaluated once, since most terms for computing the gradient analytically are already accessible as part of the cost function evaluation; time complexity being *O*(1). As shown in Fig. 4, the average computation time per iteration based on the numerical gradient is approximately (dim(***p***)+1) times of that based on the analytical, so the optimization is always much faster if the gradient is specified.

As for how the optimization progresses each iteration, the specifics depend on the algorithm. For instance, the `levenberg-marquardt` algorithm has only one conditional statement, in which a search direction–related factor is either magnified or reduced based on if a step (i.e., an iteration) succeeds in reducing the cost [43]. Consequently, the residual between the analytical and numerical gradients will not induce any difference in the optimization, unless the accumulated numerical error happens to diverge a step from success to failure. For most of the initial points we tested, the optimization process is identical between the analytical and numerical gradients (Fig. 2 and Supplementary Figs. 3–5). This also verifies our derivation of the analytical gradient.

With the `trust-region-reflective` algorithm, there is a much larger difference in the results between the gradient-specified and unspecified optimizations (Fig. 3 and Supplementary Figs. 6–10). The reason is that this algorithm not only contains a conditional statement with three branches that could separate the optimization process, but these branches are unique in terms of determining the direction of a search step: When the Hessian of the cost function is positive definite, the Newton’s method is applicable for rapid local convergence (namely a step in the Newton direction), otherwise the solution must be approached either in the direction of the steepest descent or negative curvature [42], [43]. Since the Hessian is approximated using quasi-Newton methods [42], if the gradient itself is already a numerical estimation, the Hessian approximation might not be positive definitive even if its exact value is. In this case, a Newton step is no longer possible, and neither of the alternatives converges to the solution as quickly. With the analytical gradient, however, the approximation is closer to the exact Hessian. Based on our observation so far, for every iteration, the Hessian approximated based on the analytical gradient maintains positive definite, indicating that Newton steps are likely taken throughout the optimization even with alternatives in `trust-region-reflective,` hence rapid convergence is ensured. Notably, the positive definiteness of the Hessian also allows the application of conjugate gradient methods to obtain each Newton step [42], avoiding the need for inverting the Hessian using Gaussian elimination, which is computationally less efficient.

To encapsulate, specifying the gradient in its analytical form benefits the optimization in two ways:

1. efficient computation of every iteration, and
2. rapid convergence (i.e., fewer iterations) in trust-region methods.

These two advantages grant the gradient-specified optimization much more cost descent per function count compared with the unspecified, allowing the search from many more initial points for the global solution. For instance, this rapidly convergent iteration process can be utilized for selection and mutation in genetic algorithm [46]–[48] and guarantees producing the locally optimal child without generating a large population.

With these advantages, the specification of gradient resolves most difficulties for the optimization in practice, but not all of them. Ideally, had the solver run for enough time with a sufficiently large number of iterations, there is not much need for any further conditions or constraints. However, expenditures in computation and time are always a concern in reality, and we may as well include some conditions as long as the required information is not more than we already have. To begin with, the type of obstacle as well as the number are predetermined in our method to define the vector structure of path parameters. It is of course possible to introduce a soft function that merges moment arm computation with different amount of various obstacles, so that the obstacle type and number can also be optimized. Yet we find it unnecessary at the current stage, since the cylinder is a more popular obstacle for muscle path modeling, and the obstacle number is highly related to the number of joints a muscle crosses so it can be easily predetermined.

We also constrained the 3-D coordinates of the origin and insertion points to a range (±30 mm) around the reference values; imagine the volume of a standard Rubik’s Cube. This is not essential for our method, but it is also not necessary to spend time optimizing what is already quite clear. The origin and insertion are attached to anatomical landmarks, whose relative locations on the bones are well-studied. Especially if the goal is to calibrate upper-limb muscle path, even with individual variability taken into account, the origin and insertion points should not shift too much from our selected bounds. Other parameters, particularly those of the cylinders, were tuned without any constraint, so the major complication of obstacle placement is still being tackled without the knowledge of musculoskeletal geometry (Fig. 7). In fact, our modification of the obstacle-set method has already removed complex geometric constraints between the cylinder and the points, and the optimization may proceed from any initial point (Fig. 6).

Last, there is the typical problem of local minimum, which cannot be avoided even with the optimization gradient and constraints. For this, we adopted the common strategy of multiple initial points, and for each muscle, 84–420 different initial points could be iterated in optimization. This does not eliminate the risk of local minimum, but should avoid it as much as possible. Again, it would be neat and tempting to hit the global minimum with only one shot, but our approach is more practical with the computational cost taken into account.

The performance of our method is demonstrated by replicating the muscle paths of a reference model in four conditions (Table I). Here, note that we validated the optimized parameters using non-smooth wrapping and moment arm computation, meaning that these parameters are (sub-)optimal for the original model as well. It is hence practical to implement our method for other models: For example, with some slight modifications to accommodate other wrapping strategies (not necessary if the outcome is similar), it is possible to build upon our method a user-friendly program, which takes in any desired skeleton model and moment arm–joint angle relations for muscle path calibration reproducible in OpenSim and other software.

Moreover, we need to reiterate that it is mainly the high dimensional relation between moment arm and joint configuration that makes muscle path calibration challenging, which is often neglected. The moment arm in our reference model is not a one-dimensional variable but a vector of 12 elements. For instance, the pectoralis major crosses the shoulder complex and actuates up to eight DoFs, and each of the accordant moment arms varies with changes in any one of the eight DoFs, meaning that their moment arm–joint angle relation is depicted in a 8-D space. Calibrating the moment arms of the pectoralis major is similar to matching eight pairs of 8D hypersurfaces, and it is almost impossible to accomplish manually unless four or five moment arms are neglected. In spite of that, our method succeeds in calibrating most muscles with satisfactory speed and accuracy: e.g., the three parts of the pectoralis major were each calibrated in a few seconds, and the validation errors in their 24 moment arms are all negligible. One limitation of this study is that the performance of our method is demonstrated only with artificial data generated from a model. To eliminate the concern that our optimization performance mainly benefits from the same structure shared by the parameters in the reference model and those to be calibrated, we also tested the scenario where a simpler parameter structure is configured. As shown in Table I and Supplementary Fig. 11, though both calibration speed and validation accuracy decrease compared with when the original parameter structure is configured, the calibration time is only 10.9 min and the median validation error is 0.09 mm. In reality, the path of a muscle is geometrically far more complicated than a few points and wrapping obstacles, but our results show that the optimization could still succeed with a relatively simple parameter structure. We also tested conditions with noisy data, and our method is quite robust (Table I and Supplementary Figs. 12 and 13) against errors likely larger than expected in experimental measurements. Importantly, our method is conveniently compatible with experimental data, since the only mandatory input is the relationship between moment arms and joint angles, regardless of how the data are obtained or whether the subject is human.

Another expected concern could be that, due to current technical limits, we usually do not have access to the measurements for all moment arms of a muscle, let alone their relations with all actuating DoFs. Yet this is not a problem related to our method but rather faced by all biomechanists. When lacking measurements, the moment arms will simply be assumed of certain values or relations, which is essentially what has been done in musculoskeletal modeling. Nevertheless, had such advanced measurements been available, our method is in line to provide equally advanced calibration. In fact, even if the high dimensional moment arm–joint angle relation is not completely established, our method is already usefully applicable. For example, to calibrate the Achilles tendon plantarflexion moment arm for a subject-specific model, one may use ultrasound or MRI to perform the measurement at the neutral ankle position, and then linearly scale some generic moment arm relation [49], [50] to obtain ***r***_target_(***q***) for our method. Or if one assumes some relation between moment arm and subject anthropometry, then they can also take moment arm values in the generic model as reference, scale them based on the assumed relation, and input in our method to generate a subject-specific model. In either case, the resultant muscle path will be much more accurate than when path parameters are directly scaled based on skeletal geometry.

Given these limitations, we expect our method to be examined with more models and datasets. As aforementioned, a practical application of this method would be in subject-specific modeling to recalibrate muscle path after scaling, which will be demonstrated in our future work. We are currently utilizing this method to develop a lower-limb model based on experimental data, which should offer further validation aligned with the theoretics in the current study.

It is also worth noticing that, apart from the key role in muscle path calibration, our analytically derived gradient reveals conclusively the sensitivity of moment arm to path-related parameters. The value of each element in the gradient denotes how much influence each parameter has on moment arm in a certain joint configuration, which could be of important reference in medicine. For example, in the surgery repairing rotator cuff tears, the positional error of tendon reattachment or graft insertion would induce unwanted changes in moment arms, whose magnitudes may be different depending on the direction of positional shift. The information of sensitivity could help the surgeons or manufacturers of medical devices to focus on limiting the error in the direction with potentially larger moment arm variance, so as not to compromise the biomechanical function of the operated shoulder.

Finally, we recapitulate the universal practicality of the concept we employed in the derivation of optimization gradients. A complex function may be disassembled into a product of simple modules for separate derivation, and the gradients of each module not only can be reassembled into the desired gradient composite, but may also be used as part of the gradient in other cost functions. A good example would be *∂u/∂****p*** in (4), from which we conveniently derived the gradient for a path coordinate–based cost function. This enables calibration with medical imaging data (e.g., MRI) to expand the applicability. Furthermore, it would be interesting if kinetic measurements such as joint moment can be utilized in calibration, which requires the optimization of musculotendon parameters. In a previous study, we have analytically derived the requisite gradient [12], and it can be integrated with this work, making possible joint moment–based model calibration, which is key to automated musculoskeletal modeling.

## V. Conclusion

A gradient-based method is developed for automated muscle path calibration, where path-related parameters are optimized to minimize the error in muscle moment arm. This method features specifying the gradient in its analytical form, which enables efficient computation and rapid convergence compared with using the numerical gradient, and the performance is demonstrated by fast and accurate replication of paths from a 12-DoF 42-muscle human shoulder–arm model.

We explain the derivation in detail, which first requires to smooth the cost function by replacing the conditional statements and nondifferentiable components with soft functions, and then modularize it into multiplicative factors that are easier to derive separately. This concept applies to other cost functions as well, and should be practical in the optimization of various properties in musculoskeletal models as well as many other systems.

## A. Coordinate Transformation and Partial Angular Velocity

In our mathematical derivation, there are three types of reference frames in which the coordinates of a point are expressed: world (w), joint (j), and cylinder (c), and the transformations in-between are proceeded as

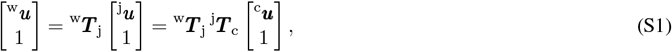

where ^w^***T*** _j_ is the transformation matrix from some specific joint frame to the world frame, and ^j^***T*** _c_ cylinder to joint. Notice the anchor points and the center(s) of the cylinder(s) may be defined with respect to different joint frames, and unless specified, j refers to the correspondent frame in which a point is fixed against joint rotation; that is, the implication of j may be different between ^j^***u***_P_, ^j^***u***_S_, and ^j^***u***_C_.

For a path segment, the computation of moment arm requires the *partial angular velocity* from the anchor points and the valid wrapping points, and it is expressed as, e.g.,

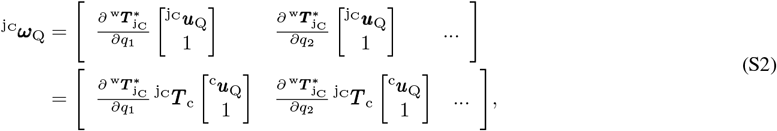

where the asterisk denotes the exclusion of the last row in the transformation matrix. The superscript refers to the joint frame in which the coordinates of the point (designated by the subscript) are expressed. Importantly, the designated point does not need to be fixed with respect to this joint frame.

## B. Verification of Smooth Modification

The smooth modification of the obstacle-set method [1] and moment arm computation is verified in a way similar to the validation process described in Subsection II-C:

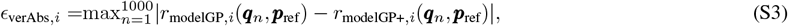

and

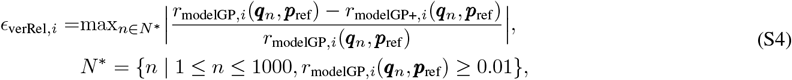

where *r*_modelGP,*i*_ is the *i*-th moment arm computed with the slightly revised obstacle-set method (Fig. 1, middle) and ***r***_modelGP+,*i*_ with the smooth form described in Subsection II-B. The input ***p***_ref_ is path-related parameters from [18], and ***q***_*n*_ is interpolated from the kinematic measurements of the shoulder and arm from 10 intransitive daily tasks (100 joint configurations per task) performed by a single subject in [41], including gesturing an OK sign, pumping fists, blocking out light from the face, saluting, pointing, gesturing a thumb-down, signaling for hitchhike, greeting, gesturing to stop, and gesturing for silence. For *ϵ*_verRel,*i*_, a joint configuration will be excluded if the correspondent |*r*_modelGP,*i*_| is below 0.01 mm (indicated by *N*^*∗*^). Moment arm is computed in a 12-DoF human shoulder–arm model [40]; see Supplementary Table I for DoFs in detail. As shown in the figures below, across 1000 joint configurations in the range of motion, the moment arm computed based on our smoothed method has negligible difference from the original value, with the maximum from 182 effective moment arms of 42 muscle paths being 0.02 mm; see Supplementary Table II for muscle paths in detail. The existing difference stems from the nature of the soft functions, rather than coding issues.

**S Fig. 1.**
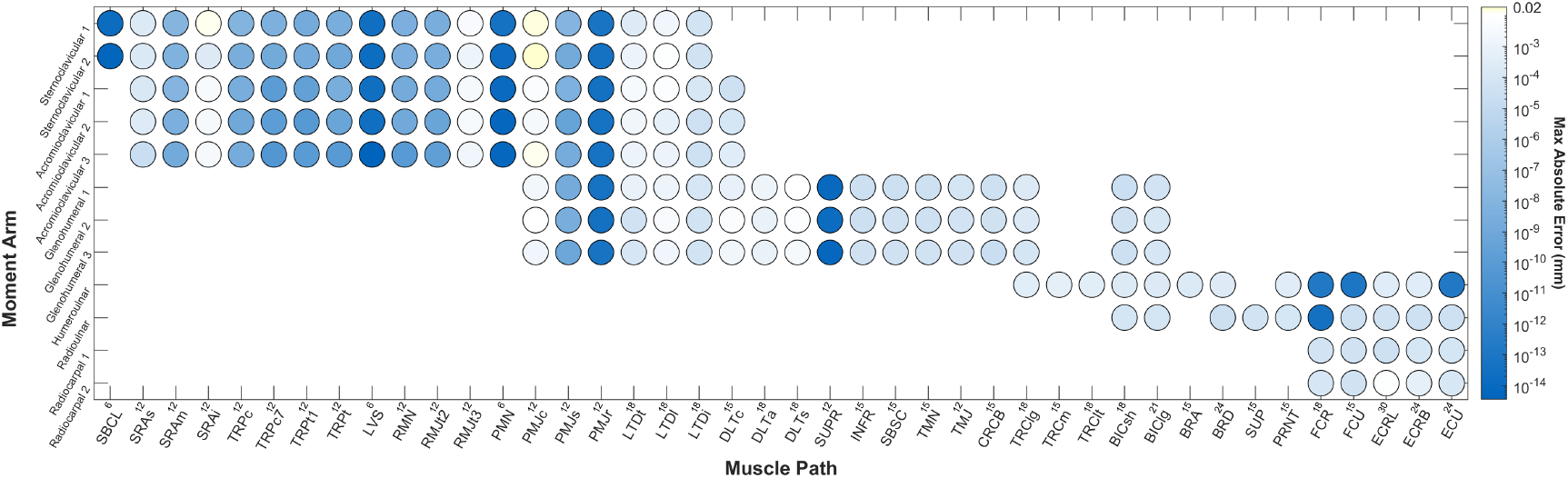
Absolute errors in 182 effective moment arms of 42 muscle paths (max value across 1000 joint configurations). The magnitude of the error is indicated by the intensity of the color. The number in the horizontal axis corresponds to the parameter number for each muscle. See Appendix D for nomenclatures in the axes.

**S Fig. 2.**
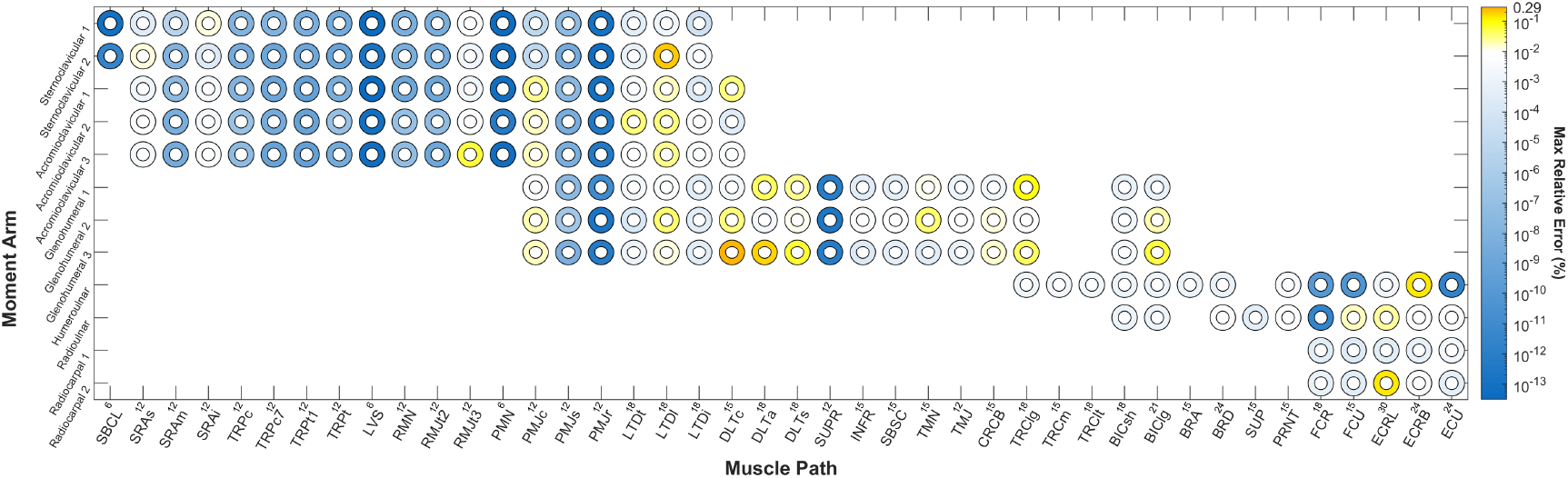
Relative errors in 182 effective moment arms of 42 muscle paths (max value across all valid joint configurations). The magnitude of the relative percentage is indicated by the intensity of the color. The number in the horizontal axis corresponds to the parameter number for each muscle. See Appendix D for nomenclatures in the axes.

## C. Derivation of Gradients

In our mathematical derivation, the gradient of a scalar variable is formatted as a row vector, while that of a vector variable is formatted as a matrix, namely the Jacobian, whose rows are the gradients of each scalar variable. Suppose for the single-cylinder case with 12 parameters, *∂R/∂****p*** is simply a 12-D row vector whose seventh element is 1 while the rest are 0, and *∂* ^j^***u***_P_*/∂****p*** is a 3*×*3 identity matrix concatenated with a 3*×*9 null matrix.

In Step 1, the coordinates of the anchor points are transformed from the joint frame to the cylinder frame, e.g.,

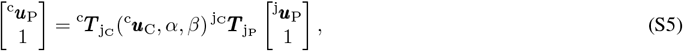

where 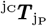 is the transformation matrix between two joint frames (dependent on ***q*** and irrelevant to ***p***), and ^c^***T*** _j_ (^c^***u***_C_, *α, β*) is the transformation matrix from the joint frame to the cylinder frame, determined by the location and orientation of the cylinder

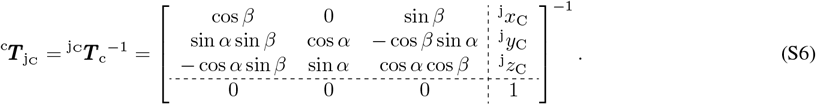

Based on the product rule, we may then easily compute the gradient, e.g.,

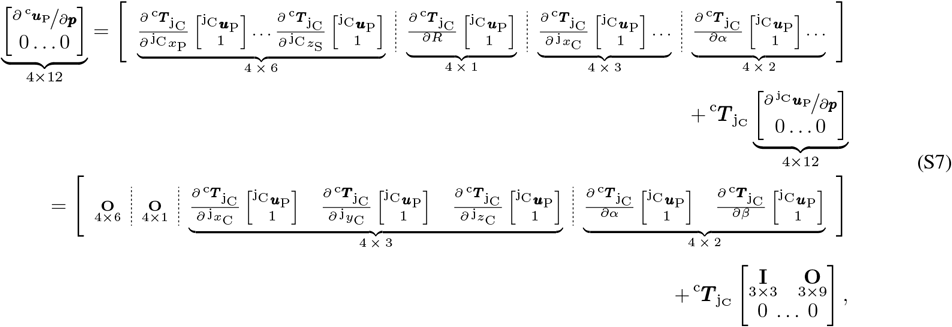

and notice the partial derivatives with respect to ***p*** are in the order of ^j^*x*_P_, ^j^*y*_P_, ^j^*z*_P_, ^j^*x*_S_, ^j^*y*_S_, ^j^*z*_S_, *R*, ^j^*x*_C_, ^j^*y*_C_, ^j^*z*_C_, *α*, and *β*.

Step 1.5 checks if the anchor points are outside the cylinder, and we utilized (8) to return adjusted coordinates as well as their gradients, e.g.,

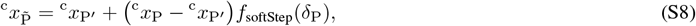

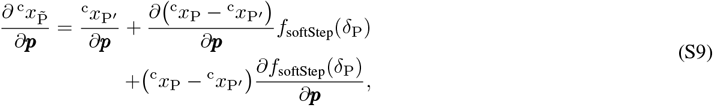

where all of the partial derivative terms can be easily derived using the chain rule. For instance,

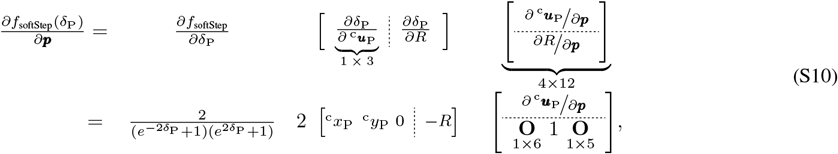

where *∂* ^c^***u***_P_*/∂****p*** is the only variable unspecified in ***p*** and may be obtained from (S7). To avoid inflating the equations, we omit the process of expanding each and every partial derivative terms in the rest of this section and focus on the format instead.

Next, with 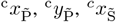, and 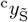 from (S8), we can derive the *x* and *y* coordinates of the wrapping points as described in [1]:

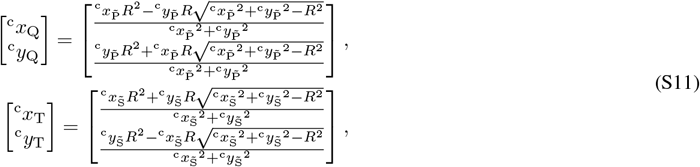

and additionally with the respective partial derivatives from (S9), we can also compute their gradients

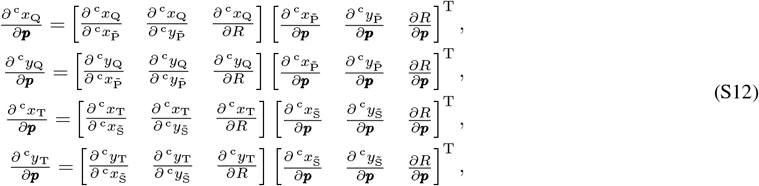

where the first terms on the right side are easy to derive from (S11), and the second terms are obtained from (S7) and (S9).

Step 2 ends with the calculation of ^c^*z*_Q_ and ^c^*z*_T_ as described in [1]:

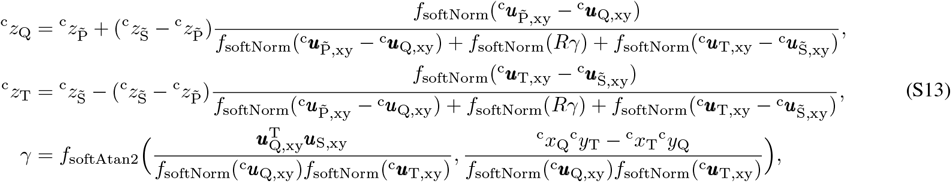

and Step 3 returns the coordinates of 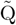 and 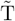 after checking the validity of the wrapping with (12). Likewise, their gradients are formatted as

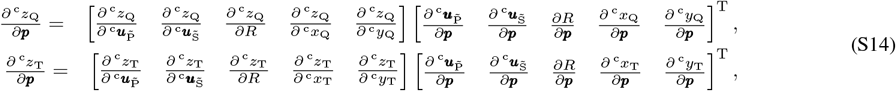

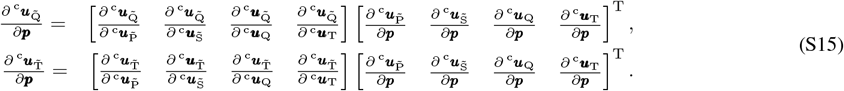

Here at the end of the obstacle-set method, we have obtained 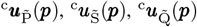 and 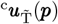 as well as their gradients. Before computing moment arm with (13), we need to utilize (S1) to transform them from their respective joint frames to the world frame.

Finally, (4) may now be extended into

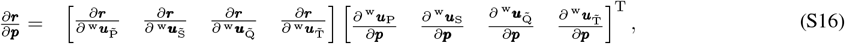

where the first term on the right side is easily derived from (13), while the second term lies in the chain of (S7), (S12), and (S12)–S15. The derivation of this compound gradient is verified as described in Subsection II-C.

**S TABLE 1.** Nomenclature for 12 DoFs in the human shoulder and arm [40].

| Joint | DoF (Moment Arm) |
| --- | --- |
| Sternoclavicular 1 | Protraction/retraction |
| Sternoclavicular 2 | Elevation/depression |
| Acromioclavicular 1 | Lateral abduction/adduction |
| Acromioclavicular 2 | Internal/external rotation |
| Acromioclavicular 3 | Posterior abduction/adduction |
| Glenohumeral 1 | Anteversion/retroversion |
| Glenohumeral 2 | Abduction/adduction |
| Glenohumeral 3 | Interior/exterior rotation |
| Humeroulnar | Flexion/extension |
| Radioulnar | Pronation/supnation |
| Radiocarpal 1 | Palmarflexion/dorsiflexion |
| Radiocarpal 2 | Radial/ulnar abduction |

**S TABLE 2.**
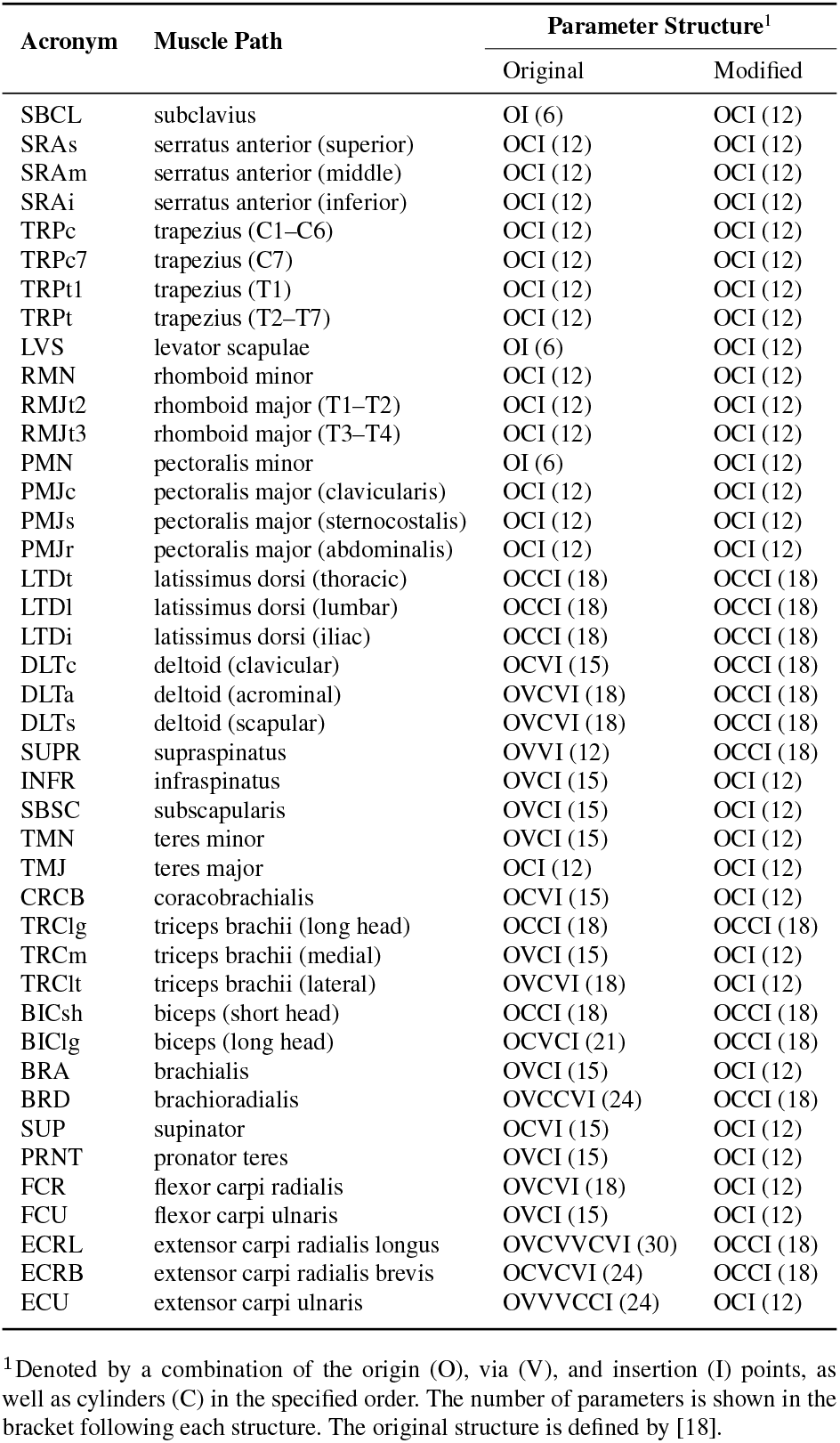
Nomenclature and parameter structure for 42 muscle paths in the human shoulder and arm.

## D. Nomenclature

## E. Supplementary Figures (Analytical vs. Numerical)

The following figures are supplements to Figs. 2 and 3, showing the cost–iteration curves for the optimization using either the algorithm of

1. levenberg-marquardt (70 initial points), or
2. trust-region-reflective (84 initial points).

The results of each initial point from the analytical (blue) and numerical (red) gradients are plotted respectively on the left and right. The mean and STD of the costs are plotted in the middle, with function count–dependent curves in the corner.

**S Fig. 3.**
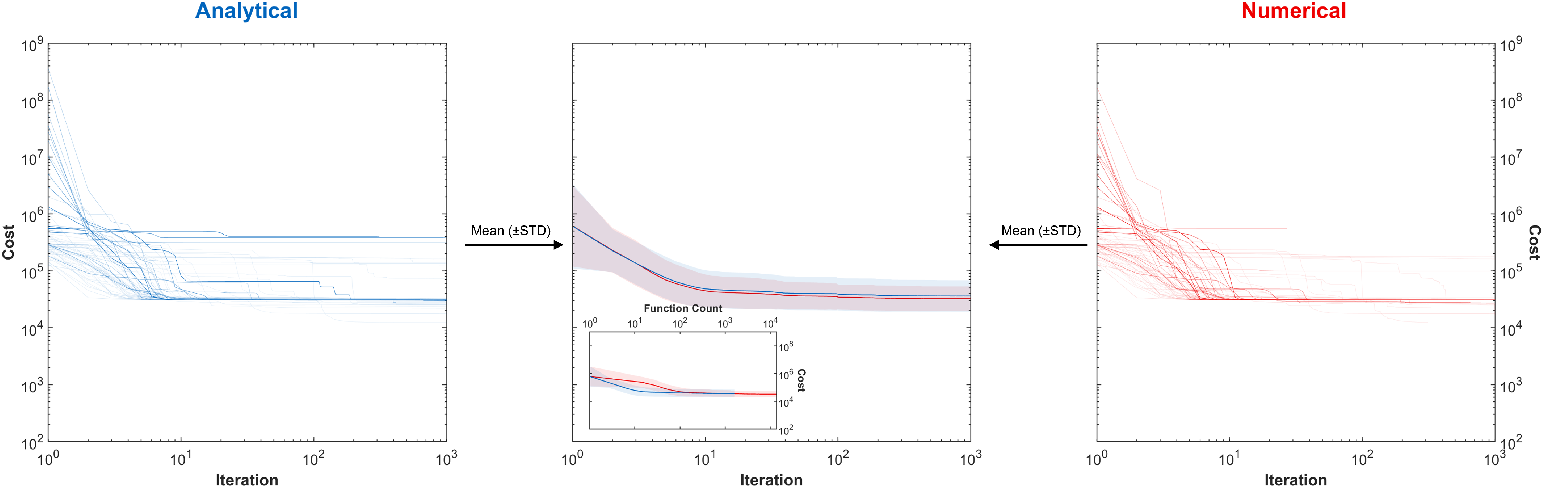
`levenberg-marquardt`, serratus anterior (superior part).

**S Fig. 4.**
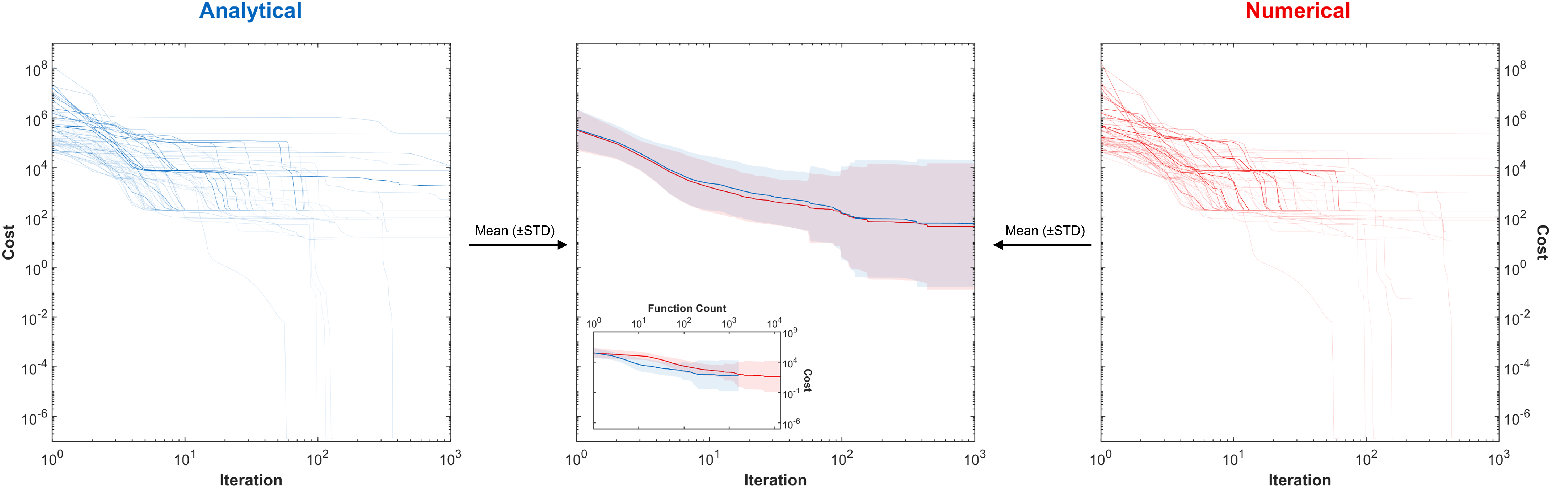
`levenberg-marquardt`, serratus anterior (middle part).

**S Fig. 5.**
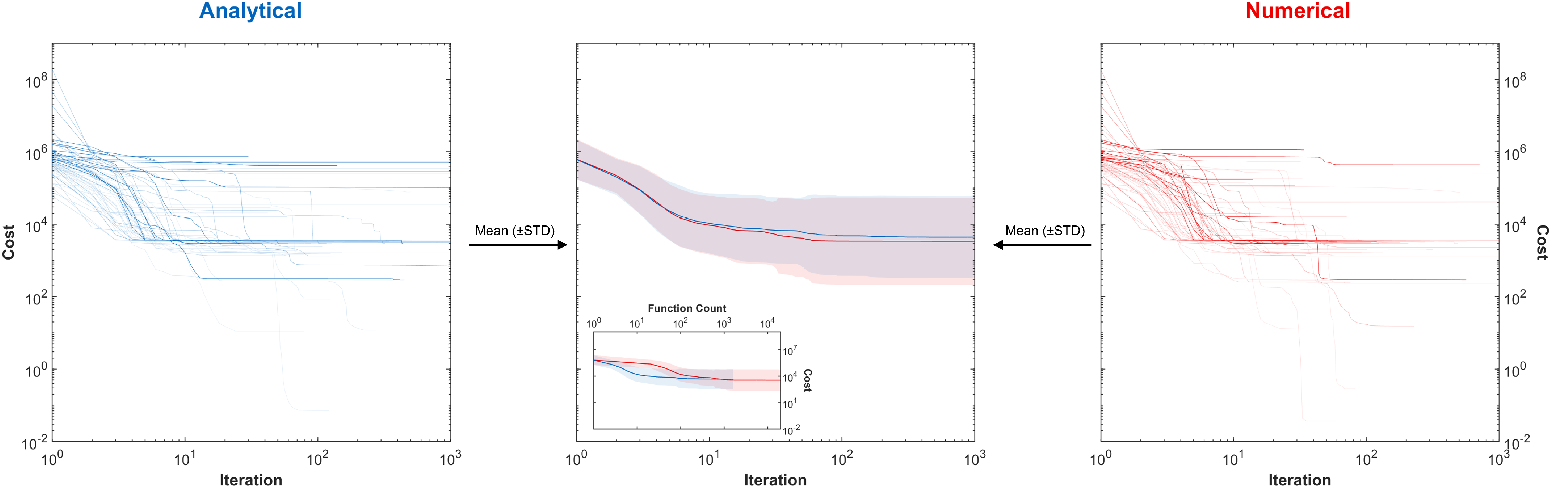
`levenberg-marquardt`, latissimus dorsi (thoracic part).

**S Fig. 6.**
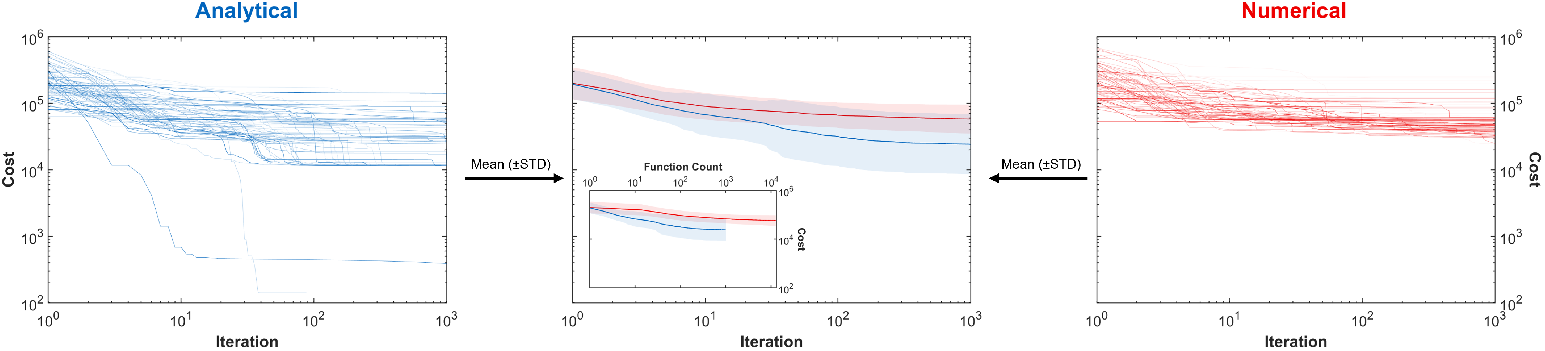
`trust-region-reflective`, serratus anterior (superior part).

**S Fig. 7.**
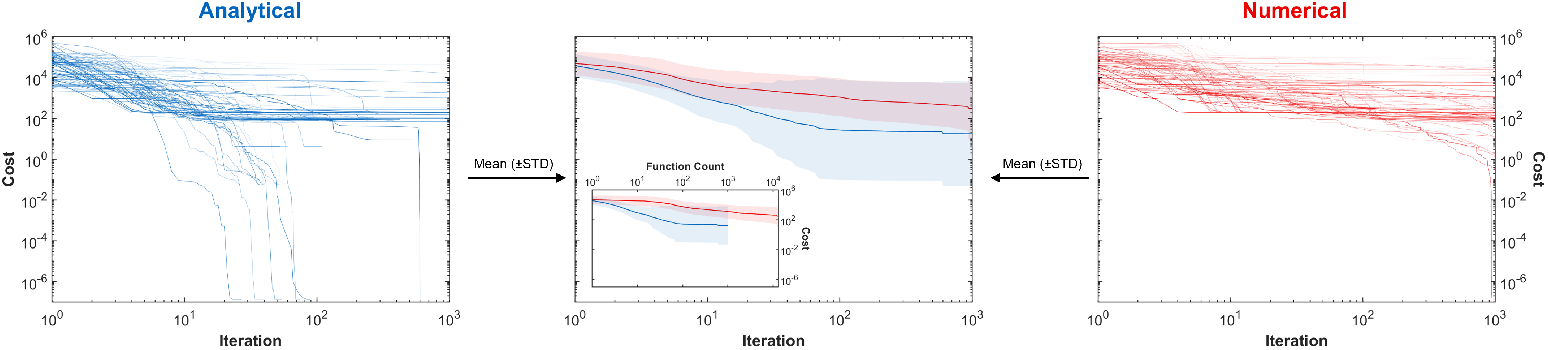
`trust-region-reflective`, serratus anterior (middle part).

**S Fig. 8.**
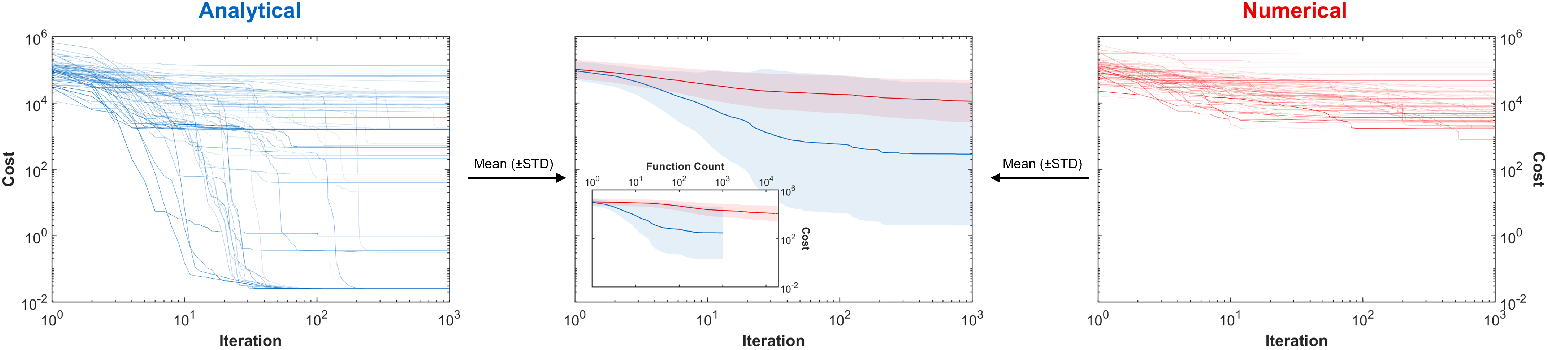
`trust-region-reflective`, latissimus dorsi (thoracic part).

**S Fig. 9.**
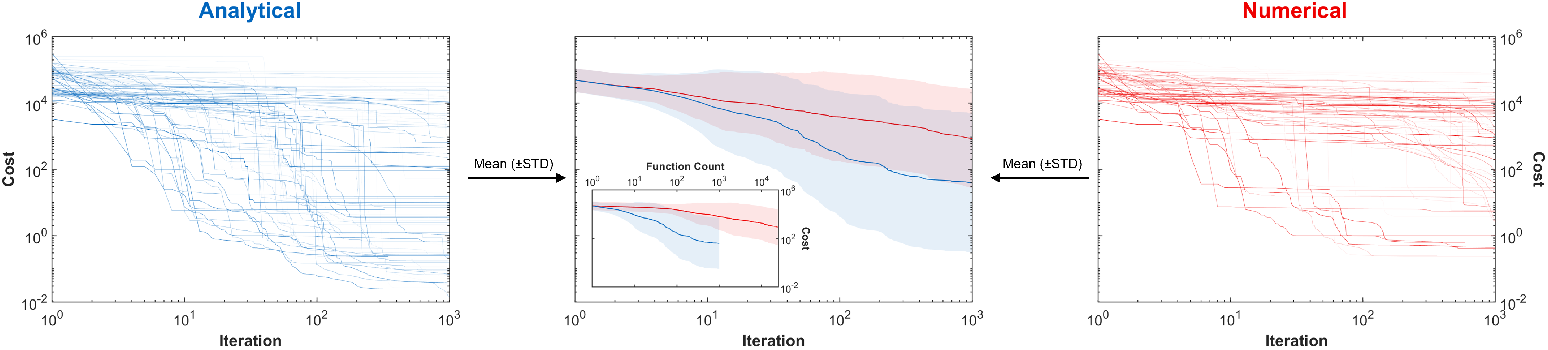
`trust-region-reflective`, extensor carpi radialis brevis.

**S Fig. 10.**
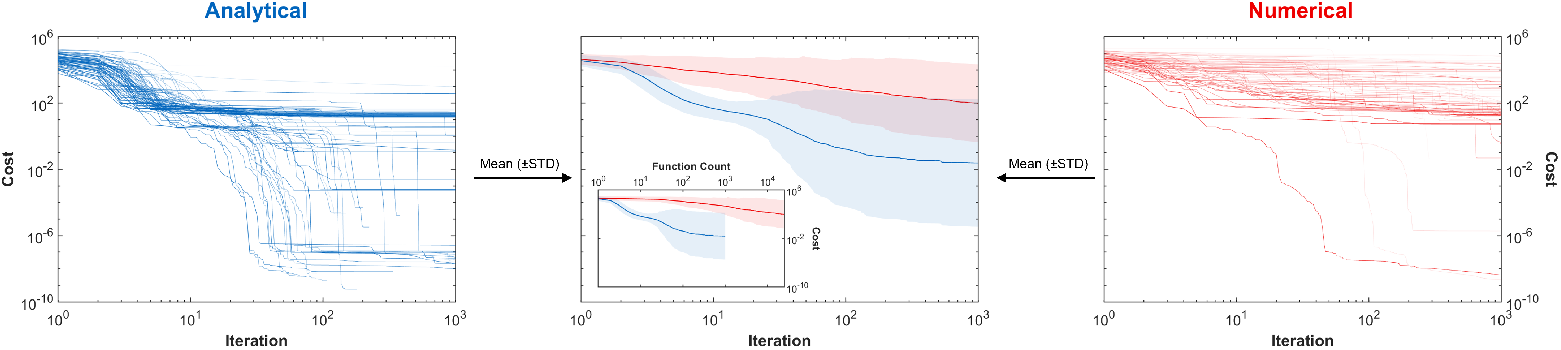
`trust-region-reflective`, extensor carpi ulnaris.

## F. Supplementary Figures (Validation Error)

The following figures are supplements to Fig. 5, showing the validation errors in 182 effective moment arms of 42 muscle paths. The magnitude of the error is indicated by the intensity of the color. The number in the horizontal axis corresponds to the parameter number for each muscle path. See Appendix D for nomenclatures in the axes.This work was supported by the Lighthouse Initiative Geriatronics by StMWi Bayern (Project X, grant no. 5140951) and the German Federal Ministry of Education and Research funding (Project AI.D, grant no. 16ME0539K).

All authors are with the Munich Institute of Robotics and Machine Intelligence (MIRMI), Technical University of Munich (TUM), Germany.

Tingli Hu and Sami Haddadin are also with the Chair of Robotics and Systems Intelligence (RSI), TUM School of Computation, Information and Technology, Germany

Ziyu Chen and David W. Franklin are also with Neuromuscular Diagnostics, TUM School of Medicine and Health, Germany. David W. Franklin is also with the Munich Data Science Institute (MDSI), TUM, Germany.

**S Fig. 11.**
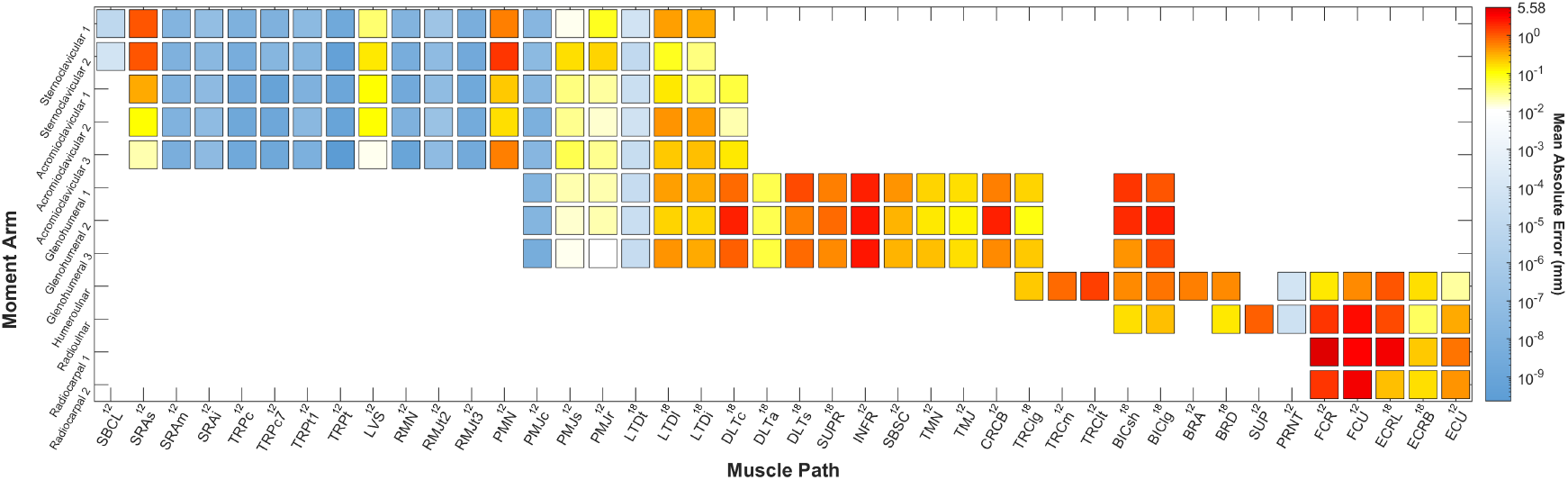
Modified parameter structure, noise-free calibration data.

**S Fig. 12.**
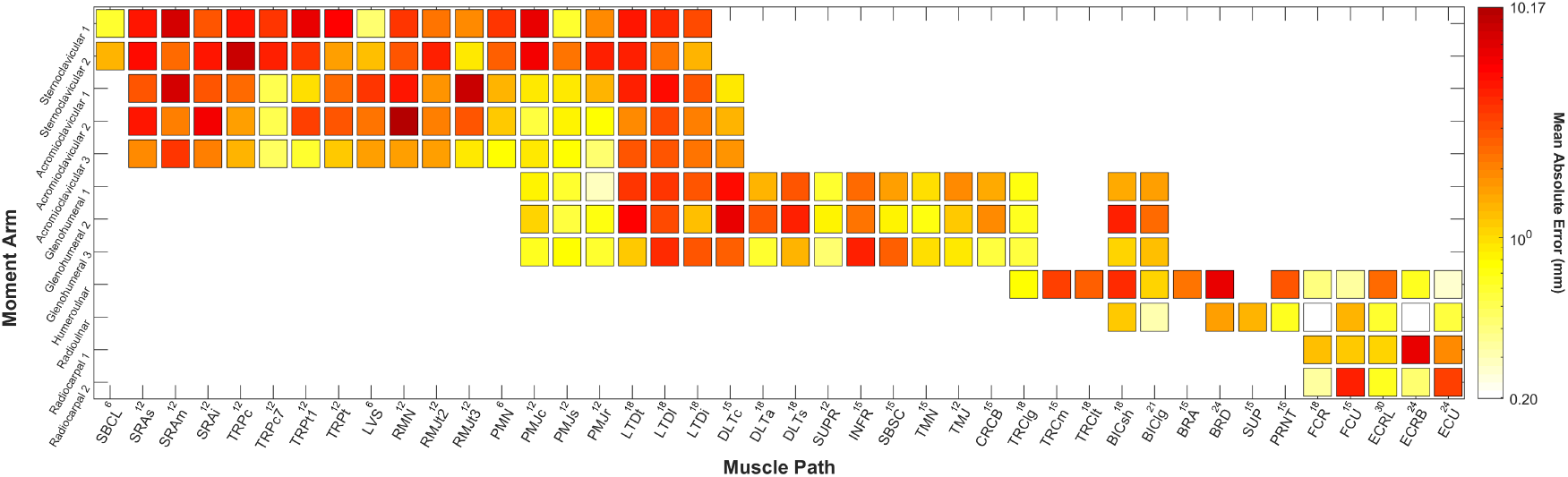
Original parameter structure, noisy calibration data.

**S Fig. 13.**
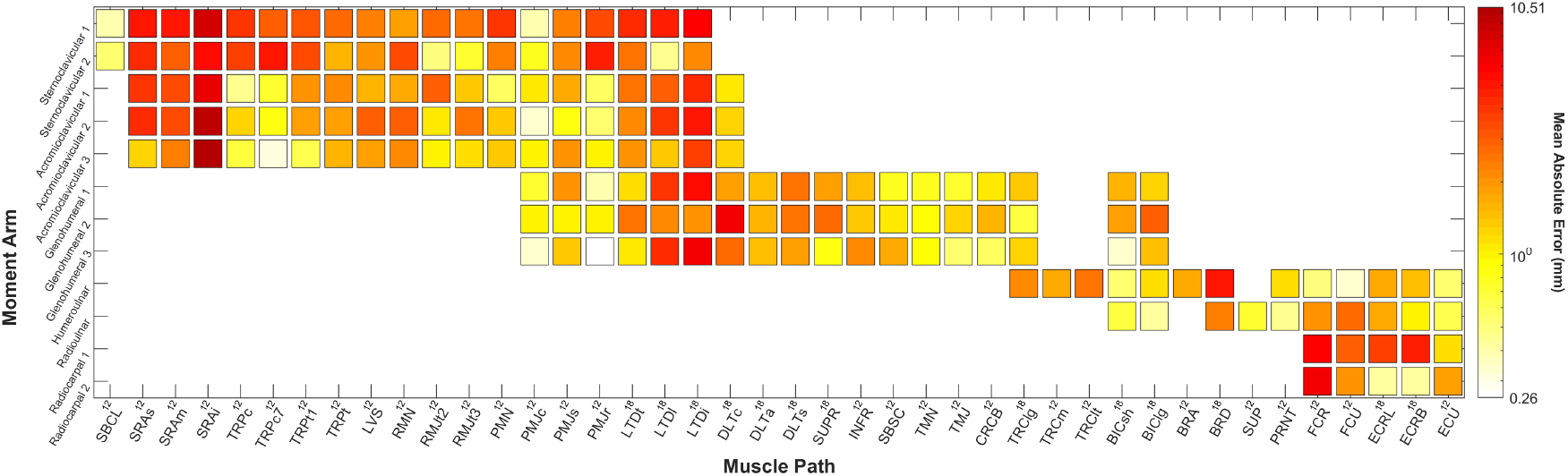
Modified parameter structure, noisy calibration data.

